# Neuromodulation leads to a burst-tonic switch in a subset of VIP neurons in mouse primary somatosensory (barrel) cortex

**DOI:** 10.1101/475061

**Authors:** Alvar Prönneke, Mirko Witte, Martin Möck, Jochen F. Staiger

**Affiliations:** Institute of Neuroanatomy, Universitätsmedizin Göttingen, Georg-August-Universität, Kreuzbergring 36, Göttingen D-37075, Germany

**Keywords:** Acetylcholine, Barrel Cortex, Burst Spiking, GABAergic Interneurons, Serotonin, Vasoactive Intestinal Polypeptide

## Abstract

Neocortical GABAergic interneurons expressing vasoactive intestinal polypeptide (VIP) contribute to sensory processing, sensorimotor integration and behavioral control. In contrast to other major subpopulations of GABAergic interneurons, VIP neurons show a remarkable diversity. Studying morphological and electrophysiological properties of VIP cells, we found a peculiar group of neurons in layer II/III of mouse primary somatosensory (barrel) cortex, which showed a highly dynamic burst firing behavior at resting membrane potential that switched to tonic mode at depolarized membrane potentials. Furthermore, we demonstrate that burst firing depends on T-type calcium channels. The burst-tonic switch could be induced by acetylcholine and serotonin. Acetylcholine mediated a depolarization via nicotinic receptors whereas serotonin evoked a biphasic depolarization via ionotropic and metabotropic receptors in 48% of the population and a purely monophasic depolarization via metabotropic receptors in the remaining cells. These data disclose an electrophysiologically-defined subpopulation of VIP neurons that via neuromodulator-induced changes in firing behavior is likely to regulate the state of cortical circuits in a profound manner.

## Introduction

In the neocortex, excitation is controlled and shaped by inhibitory interneurons, which are also directly involved in sensory information processing and working memory (Isaacson and Scanziani 2011; Harris and Mrsic-Flogel 2013; Karnani et al. 2014; Kepecs and Fishell 2014; Tremblay et al. 2016; Kamigaki and Dan 2017). Today, three major subclasses of inhibitory interneurons are distinguished by the expression of certain marker proteins. These are parvalbumin (PV), somatostatin (Sst), and serotonin receptor 3a (5HT_3a_R) expressing interneurons, from the latter about 30% also express vasoactive intestinal polypeptide (VIP) (Rudy et al. 2011; Pfeffer et al. 2013; Staiger et al. 2015; Tremblay et al. 2016; Feldmeyer et al. 2017). However, each of these major subclasses consists of several distinct cell types, which are commonly believed to be only safely distinguished by multimodal characterization (DeFelipe et al. 2013; Zeng and Sanes 2017).

Current research focusses on VIP-expressing interneurons because they shape excitation by a distinct disinhibitory circuit motif that seems to be relevant during many aspects of behavior (Lee et al. 2013; Pi et al. 2013; Fu et al. 2014; Pinto and Dan 2015). The underlying circuitry was shown to be VIP neurons inhibiting Sst neurons, which results in a decrease of inhibition in excitatory cells, i.e. disinhibition. This was recently confirmed by a multitude of physiological studies (Caputi et al. 2009; Pfeffer et al. 2013; Jiang et al. 2015; Karnani et al. 2016; Walker et al. 2016). However, the hypothesis that VIP neurons exclusively target other inhibitory interneurons was already difficult to explain on the basis of previous data (Gonchar et al. 2007; Lee et al. 2013; Pfeffer et al. 2013) and certainly does not hold in the light of recent evidence that shows VIP synapses on GABAergic as well as non-GABAergic targets (Zhou et al. 2017). This morphological data goes well along with recent physiological and behavioral studies that imply that VIP neurons sculpt neuronal ensembles to mediate a certain behavior by direct inhibition as well as disinhibition (Pakan et al. 2016; Garcia-Junco-Clemente et al. 2017; Kamigaki and Dan 2017; Kuchibhotla et al. 2017).

An important question that arises now is how a single population of neurons allows for such diversity in connectivity? One answer can be found by characterizing in detail electrophysiological and morphological features of VIP neurons to disclose putative cell types that can be studied further for their connectional and behavioral properties, as to firmly establish a cell type (Zeng and Sanes 2017). We recently showed that at least two coarse populations of VIP neurons can be distinguished, a layer II/III versus a layer IV-VI population, which however consisted of at least partly inhomogeneous cells (Prönneke et al. 2015). The above described diversity of VIP neurons might be reflected by their rich array of specific firing patterns and morphologies. Besides continuous adapting firing patterns, irregular spiking and burst firing have been described in various studies (Kawaguchi and Kubota 1997; Cauli et al. 2000; Prönneke et al. 2015). Interestingly, burst firing, an especially powerful mode of neuronal output (Lisman 1997; Bean 2007; Cheng-yu et al. 2009), has so far been described only for layer II/III VIP neurons of the barrel and frontal cortex in mice (Kawaguchi and Kubota 1997; Prönneke et al. 2015). Burst firing is probably best known for thalamic relay neurons, together with their ability to switch to tonic mode upon depolarization. Burst versus tonic mode firing has been related to the two major behavioral states that the cortex can be found in, namely sleep and wakefulness (McCormick and Bal 1997; Sherman 2001). However, recent studies suggested that burst firing might have a more fine-grained functionality (Swadlow and Gusev 2001), which can be tuned by neuromodulation (McCormick and Bal 1997).

VIP neurons are well known for their reliable responses to neuromodulators. They are depolarized by acetylcholine (ACh) via nicotinic non-alpha7 receptors (Arroyo et al. 2012; Poorthuis et al. 2014; Koukouli et al. 2017), whereas their responsiveness to other neuromodulators like serotonin is still poorly understood (Ferezou et al. 2002; Munoz and Rudy 2014). This is surprising for the latter since VIP neurons form a major subpopulation of 5HT_3a_R-expressing interneurons (Tremblay et al. 2016). Thus, action potential firing properties, being differentially recruited into neuronal ensembles driven by state-dependent cortical functions, could be a key to better understand the roles that VIP neurons play in cortical circuitry.

In this study, we present data derived from the detailed analysis of intrinsic electrophysiological properties of individual VIP neurons from layer II/III of mouse barrel cortex. Firstly, burst spiking (BS) VIP neurons change their firing pattern to tonic upon depolarization. Secondly, only half of all layer II/ III VIP neurons display 5HT_3a_R-mediated responses but all are depolarized by metabotropic 5HT_2_R. Furthermore, we confirm that the depolarization induced by ACh is mediated exclusively by nicotinic receptors. Thirdly, the depolarization induced by either, ACh or 5HT, is sufficient to change the firing pattern from burst to tonic in BS VIP neurons.

## Material and Methods

### Animals

Homozygous Vip-ires-cre (VIPtm1(cre)Zjh, The Jackson Laboratory, Bar Harbor, USA) mice were crossed with homozygous Ai9 mice ((Madisen et al. 2010); floxed tdTomato mice: B6.Cg-Gt(ROSA)26Sortm9(CAG-tdTomato)Hze/J, floxed YFP mice: B6.Cg-Gt(ROSA)26Sortm3(CAG-EYFP)Hze/J, The Jackson Laboratory, Bar Harbor, USA). In the present study we used offspring (21-36 days, average 28.4±3.6) heterozygous for VIPcre/tdTomato or VIPcre/YFP from the breeding facility of the University Medical Center Göttingen (Göttingen, Germany). These mice are highly sensitive and specific (~99%) for VIP expressing interneurons (Taniguchi et al. 2011; Prönneke et al. 2015). All experimental procedures were performed in accordance with German laws on animal research (TierSchG und TierSchVersV 2013).

### In vitro electrophysiology

VIPcre/tdTomato mice were deeply anesthetized with isoflurane and decapitated. Thalamocortical slices (300 µm; (Porter et al. 2001)) containing the primary somatosensory (barrel) cortex were cut using a vibratome (Leica VT1200S, Wetzlar, Germany). The cold (4°C) cutting solution contained (in mM): 75 sucrose, 87 NaCl, 2.5 KCL, 0.5 CaCl2, 7.0 MgCl2, 26 NaHCO3, 1.25 NaH2PO4 and 10 glucose, and was continuously equilibrated with 95% O2 and 5% CO2, pH 7.4. Slices were incubated for 0.5-1h at 34°C prior to recording in extracellular solution (ACSF) of the following composition (in mM): 125 NaCl, 2.5 KCL, 2 CaCl2, 1 MgCl2, 26 NaHCO3, 1.25 NaH2PO4 and 25 glucose, pH 7.4 when equilibrated with 95% O2 and 5% CO2. Slices were transferred to a fixed-stage recording chamber in an upright microscope (Axio Examiner, Zeiss, Germany) and continuously perfused at a rate of 2 ml/min with ACSF. All experiments were performed at 32°C. The barrel field was visualized at low magnification (2.5x) under bright-field illumination. Target neurons were identified by tdTomato or YFP fluorescence using a 40x water immersion objective (40x/0.75W, Olympus, Germany). For whole-cell patch-clamp recordings, filamented borosilicate glass capillaries (Science Products, Hofheim, Germany) of 5-8 MΩ resistances were filled with (in mM): 135 K-gluconate, 5 KCL, 10 HEPES, 0.5 EGTA, 4 Mg-ATP, 0.3 Na-GTP, 10 phosphocreatin phosphate and 0.3-0.5% biocytin. If not stated otherwise, membrane potentials were not corrected for a liquid junction potential of ~16 mV. Membrane potentials and currents were recorded using a SEC 05L amplifier (npi electronics, Tamm, Germany) in discontinuous current-clamp mode or voltage-clamp mode with a switching frequency of 50 kHz, filtered at 3 kHz, and digitized at 10-25 kHz using a CED Power1401 (CED Limited, Cambridge, England). Access resistance was monitored and compensated if changes appeared. Recordings during which the access resistance could not be compensated were discarded. Data were collected, stored and analyzed with Signal 5 software (CED Limited, Cambridge, England). Passive and active properties of neurons were determined immediately after establishing whole-cell configuration by applying one second lasting hyperpolarizing or depolarizing rectangular current pulses of varying strength at resting membrane potential.

### Pharmacological experiments

Neurons were exposed to pharmacological agents either by bath application or short and local pressure application. To explore the mechanism underlying burst firing and the burst-tonic switch, T-type calcium channels were blocked by bath application of 3,5-dichloro-N-[1−(2,2-dimethyl-tetrahydro-pyran-4-ylmethyl)-4-fluoro-piperidin-4-ylmethyl]-benzamide (TTA-P2, 20 µM; Tocris, Wiesbaden, Germany). In a second series of experiments, Tetrodotoxin (TTX, 0.5 µM; Tocris, Wiesbaden, Germany) was added to the bath prior and together with TTA-P2 in order to isolate the T-current mediated membrane potential response in current clamp. The ionic basis of rebound depolarizations was studied using 4-Ethylphenylamino-1,2-dimethyl-6-methylaminopyrimidinium chloride (ZD7288, 10 µM; Tocris, Wiesbaden, Germany), a potent HCN channel antagonist. Bath application of neuromodulators was monitored by continuously recording in current-clamp mode for 5 to 10 minutes. To control for changes in input resistance, every 6 seconds a hyperpolarizing stimulus of −10 pA lasting 200 ms was applied. Recovery from bath application was monitored for up to 30 minutes using the same stimulation protocol. Bath-applied neuromodulators were: acetylcholine (ACh, 40 µM; Sigma, Deisenhofen, Germany), and serotonin (5HT, 5 µM; Tocris, Wiesbaden, Germany). Bath-applied receptor antagonists were: mecamylamine (non-alpha7 nicotinic ACh receptor antagonist, 100 µM; Biotrend, Zürich, Switzerland), tropisetron (5HT_3a_R antagonist, 10 nM; Biotrend, Zürich, Switzerland), and cinanserin (5HT_2a_ and 5HT_2c_ receptor antagonist, 400 µM; Tocris, Wiesbaden, Germany). Pressure application of ACh (100 µM) and 5HT (200 µM) was performed by placing a micropipette at a distance of 15-20 µm to the soma of the recorded neuron and applying short pressure pulses (50 ms, 100 ms, or 30 s) of 7 psi using a pressure application system (Toohey Spritzer, Toohey Company, Fairfield, NJ). Short and local pressure experiments were recorded in voltage clamp with the membrane potential of neurons being clamped to −65 mV. Note that when cinanserin was bath-applied it was also added to the solution in the pressure application pipette.

### Staining of biocytin-filled cells

Staining of biocytin-filled cells has previously been described in detail (Staiger et al. 2004; Staiger et al. 2014). In brief, after recording slices were fixed in 4% formaldehyde (in PB) at 4°C for 12 to 20 hours. To stop fixation, the tissue was rinsed extensively with PB including an intermediate step with 1% H_2_O_2_ (in PB) to block endogenous peroxidase activity. Next, slices were incubated in a cryoprotectant (25% saccharose, 10% glycerol in 0.01 M PB) for 1h. They were freeze–thawed three times over liquid nitrogen. After three rinses in PB, the slices were incubated overnight with Avidin-Biotin Complex (ABC; 1:200; Vector, Burlingame, CA) at 4°C. Afterwards, slices were preincubated with 1 mg/ml 3,3′diaminobenzidine (DAB; Sigma, Deisenhofen, Germany) in PB for 10 min and the peroxidase reaction was started by adding 0.01% H_2_O_2_. The reaction was stopped by rinsing with PB. To intensify the reaction product, sections were incubated in 1.4% silver nitrate at 56°C for 30 min, followed by 0.2% gold chloride at room temperature for 10 min, and fixed with 5% sodium thiosulfate for 5 min. The barrel field was visualized by standard cytochrome oxidase histochemistry (Wong-Riley and Welt 1980).

### Reconstruction of biocytin-filled neurons

Neurons with consistently intense staining of neurites and no obvious truncation of major processes were reconstructed using live digital images acquired by a digital camera (CX9000, MBF Bioscience, Colchester, VT) mounted on a microscope (Eclipse 80i, Nikon, Ratingen, Germany) with a 63x oil-immersion objective (NA = 1.4) and connected to a computer running Neurolucida (MBF Bioscience, Colchester, VT). Dendritic processes were distinguished from axonal structures by their diameter, fine structure, and branching pattern (Prönneke et al. 2015).

### Analysis of electrophysiological data

#### Basic characterization

Electrophysiological data were analyzed using custom written scripts for Signal 5. Most of the passive properties were analyzed using averages of ten responses to a hyperpolarizing current pulse of 10 pA. The membrane time constant was determined by fitting an exponential to the averaged membrane potential response (f(x) = Ae^-x/τ^ + B; A = maximum amplitude, B = membrane potential at maximum amplitude of response; R^2^ > 0.9 in all analyzed cells). Input resistance was determined according to Ohm’s law using the maximum amplitude of the membrane potential response and the amplitude of the injected current. The sag index is an indicator of slow inward rectification. It was calculated by comparing the input resistance at the maximum amplitude to that at steady state (averaged membrane potential toward the end of the stimulus). Differences are given as percentages. Fast inward rectification, i.e. at the initial maximal membrane potential response, (rectification index, RI) was measured by comparing the maximal membrane potential changes as responses to current pulses of 10 pA to those of 100 pA. Linearity (response to 10 pA is 10% of that to 100 pA) suggests that the deflection depends exclusively on passive membrane properties. An increase in the percentage indicates involvement of fast inward rectification. For clarity, 10 was subtracted from the resulting values, thus no fast inward rectification is set to 0. Firing threshold, time to peak, half width, amplitude (measured from firing threshold to peak) of action potentials were analyzed using action potentials evoked at rheobase. To measure the firing threshold, we determined the point in time, within a time window of 1 ms before the spike peak, at which the slope of the membrane potential response dropped to 10% of the average slope within that time window. The membrane potential at this point in time was taken as firing threshold. The amplitude of afterhyperpolarizations (AHP) was determined by measuring the difference in voltage from firing threshold to maximum deflection of the repolarization. The time to peak of AHP was measured from the time point the repolarization of the action potential crossed the firing threshold to the maximum amplitude of the AHP. The distance to the pial surface of each recorded neuron was measured in images capturing the position of the tip of the recording electrode following experiments.

#### Analysis of the frequency range

To analyze the frequency spectra of VIP neurons, all instantaneous frequencies (IFF; reciprocal of inter-spike intervals) were pooled for each neuron and binned in 10 Hz steps from 10 Hz to 400 Hz and plotted as percent in a cumulative manner. Based on this, the dynamic frequency range (DFR) was defined as 5% to 98% of all instantaneous frequencies found in corresponding frequency bins. Also, the first frequency bin in which more than 50% of all frequencies are found is given in the results (Figures 2B + C). For a direct comparison of burst and tonic mode of BS VIP neurons, the non-cumulative percentile distribution of IFFs was tested for significance in each bin, and plotted in Figure 2D.

#### Analysis of pharmacological experiments

Effects of pharmacological treatments were analyzed using custom written scripts for Signal 5. The change in frequency spectra induced by blockade of T-type calcium channels was analyzed as described above. To quantify the magnitude of membrane potential responses evoked by t currents (isolated by TTX application), we calculated the amplitude and integral of depolarizing humps, relative to steady-state levels, before and during the application of TTA-P2. The effects of h currents were quantified by calculating the sag index as described above and the amplitude of the rebound depolarization, relative to baseline membrane potential, following a current stimulus of −100 pA before and during the application of ZD7288. Steady-state membrane potential changes induced by bath-applied neuromodulators were analyzed by measuring the membrane potential as the average of 10 epochs of 100 ms each just prior to the hyperpolarizing current stimuli during (a) the first 60 s of the recording, representing baseline, and (b) the last 60 s of the recoding. The amount of recovery of the membrane potential was analyzed in the same manner. To analyze the magnitude of membrane currents evoked by local pressure application, we measured the amplitude of evoked currents relative to baseline holding current levels.

### Analysis of morphological data

Properties of reconstructed neurons were quantified with Neurolucida Explorer (MBF Bioscience, Colchester, VT). Data were not corrected for tissue shrinkage. However, from several measurements, the shrinkage was determined to be around 10–15 % in the x-/y-axes and 40–60 % in the z axis. To determine the distribution of neuronal processes and superimpose multiple neurons, reconstructed neurons were registered into one file using defined layer borders as a reference (Prönneke et al. 2015), somata aligned at the same horizontal level, and neurites of individual neurons plotted as a binary image. Vertical and horizontal density of dendrites and axon were determined by calculating the vertical and horizontal average pixel density of these binary images using ImageJ (Rasband, W.S., ImageJ, U. S. National Institutes of Health, Bethesda, Maryland, USA, https://imagej.nih.gov/ij/, 1997-2016.). Furthermore, binary images were also generated from superimposing neurites of multiple neurons and filtered using a Gaussian filter with a comparable radial sigma (20 at 300 dpi) for all structures and a color look-up-table ranging from cold (blue and green for white to light gray) to warm colors (yellow and red for dark gray to black) was applied to the resulting grayscale image. This was then merged with the original black and white image and resulted in heat-maps visualizing areas of highest density of dendritic and axonal trees.

### Statistical tests

For statistical comparisons, data were tested for normality (Shapiro-Wilk test) and equal variance. If both passed, a one-way student t-test was used. If one or both failed, a Mann-Whitney rank sum test was used. P-values were adjusted to preclude alpha-inflation (Bonferroni) when necessary. For any multiple group comparison one way analysis of variance (ANOVA) was used. When significant differences were found and normality or equal variance tests passed, a post-hoc Holm-Sidak method as an all pairwise multiple comparison procedure was applied. If normality or equal variance tests failed, ANOVA was based on ranks with post-hoc Tukey or Dunn’s method as an all pairwise multiple comparison procedure. For comparison of proportions, we used a Chi-square test with Yates continuity correction. All statistical tests were performed with SigmaPlot (Version 13.0, Systat Software, Inc., Erkrath, Germany). Values are given as mean±SD if not indicated otherwise.

## Results

VIP neurons in layer II/III of the mouse barrel cortex display a great diversity in firing patterns (Kawaguchi and Kubota 1997; Cauli et al. 2000; Prönneke et al. 2015). A subset of these neurons show high frequency discharges of APs, a firing behavior often described as bursting. However, bursting is an ambiguous term used for a multitude of high-frequency events. In this study, we define bursting as firing of multiple APs with high frequency only if it already occurs at minimal current stimulation (rheobase). Using this classification, we identified 20% (55 of 269) of all recorded VIP neurons as burst spiking (BS) VIP neurons in layer II/III of the mouse barrel cortex. In layers IV-VI, however, we never observed BS VIP neurons as of yet (0 of 121).

### Electrophysiological properties of BS VIP neurons at resting membrane potential

Just below firing threshold, BS VIP neurons show a depolarizing hump during the initial phase of current stimulation (see grey traces in Figure 1A) which, on average, peaked at −45.3±3.5 mV (Figure S2A). Rheobase stimulation elicits an additional, steeper depolarizing envelope on which eventually multiple APs with short inter-spike intervals (ISIs) are evoked (see black traces in Figure 1A at rheobase stimulation). The magnitude of both transient depolarizations varied between individual BS VIP neurons and so did the number of APs comprising bursts (compare individual neurons in Figure 1A). The bursts consisted of 2 to 6 APs of variable intrinsic firing frequency. For a detailed description, we determined the instantaneous firing frequencies (IFF, see Material and Methods for details) for all ISIs in bursts at rheobase stimulation. Including all individual BS VIP neurons, irrespective of the number of APs, the average intra-burst firing frequency was 176±64 Hz. However, the IFF depends on the number of APs within a burst: bursts consisting of 2 APs had the lowest average IFF of 111±28 Hz in contrast to those comprising 6 APs with an average IFF of 217±71 Hz (for a more detailed analysis see Figure S1).

**Figure 1:**
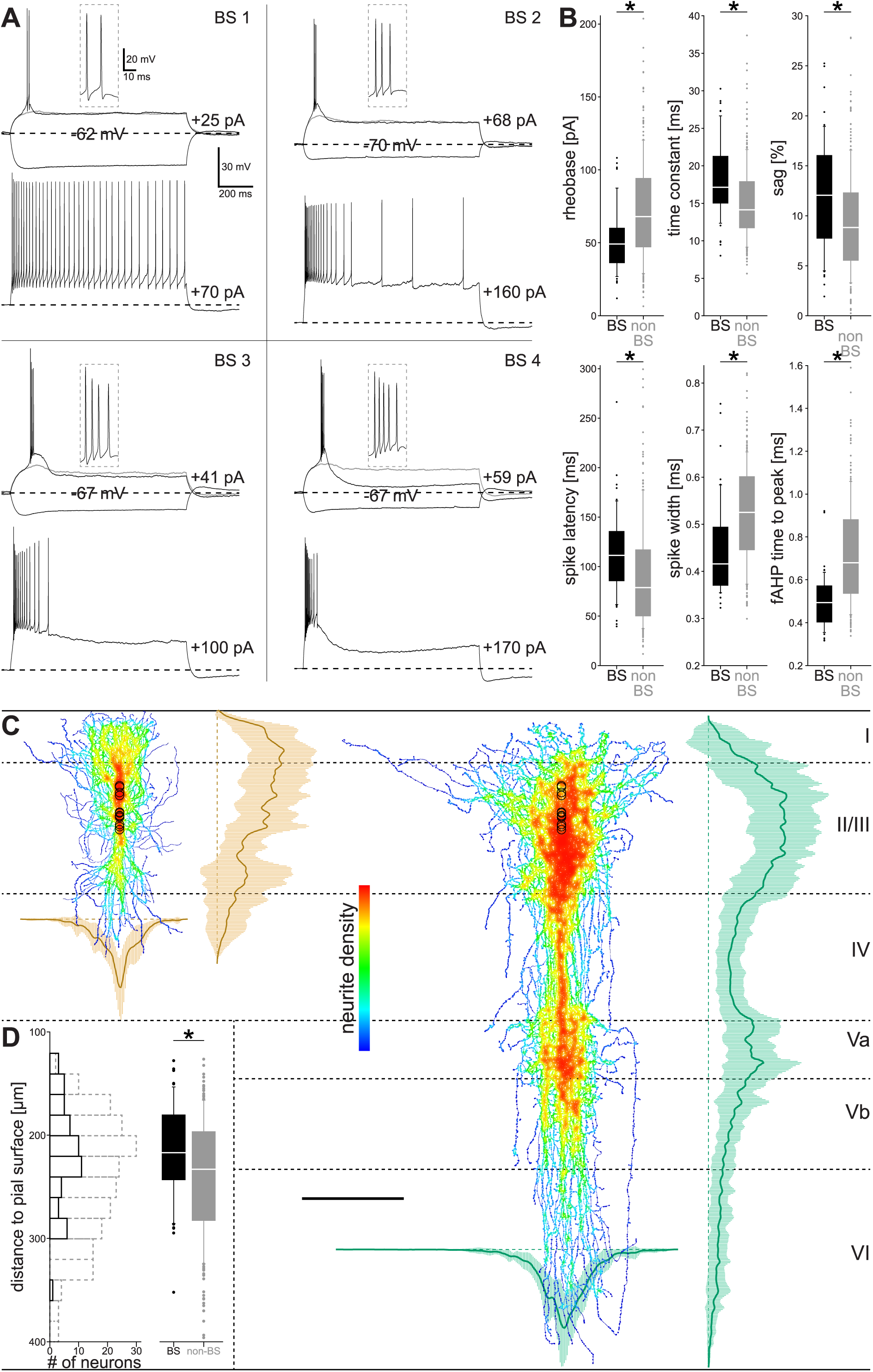
Burst spiking VIP neurons are a subpopulation of layer II/III VIP neurons In layer II/III of mouse barrel cortex, ~20% of all VIP neurons fired a high-frequency burst of action potentials at rheobase stimulation. (A) Membrane potential responses of four individual BS VIP neurons (BS 1- 4) to rectangular current stimulations (1 s) of different amplitudes (i-iii shown at the top, iv at the bottom): (i) −50 pA (black), (ii) just-subthreshold (grey), (iii) rheobase stimulation (black, current intensity indicated to the right), insets show action potential at higher temporal resolution, and (iv) clearly suprathreshold (current intensity indicated to the right). Stippled lines indicate RMP. (B) Rheobase, membrane time constant, sag index, action potential latency and width, as well as the time to peak of AHPs of BS VIP neurons (n = 55) significantly differed to non-BS VIP neurons (n = 214). (C) The morphology of the neurites of layer II/III BS VIP neurons. Dendritic trees are largely restricted to layers I and II/III and axons spread throughout all layers but preferred layers II/III and Va. The figure shows a superimposition of 12 vertically aligned somato-dendritic (left) and axonal reconstructions (right), the positions of the somata are marked by black circles. The density of neurites is indicated by a heat map ranging from warm (most dense) to cold colors (least dense). Plots show the density of dendrites (orange; average as a solid line ± SD as shaded area) and axon (in green); vertical distribution on the right of the heat maps, and horizontal distribution beneath the heat maps. Roman numerals depict layers; scale bar = 200 µm). (D) Soma location of BS VIP neurons within layer II/III (n = 55) relative to the pial surface plotted as histogram (bin size = 10 µm) and box plots. BS VIP neuron somata were found more abundantly in upper layer II/III in comparison to non-BS VIP neurons (n = 214). For electrophysiological recordings and morphology of non-BS VIP neurons see Prönneke et al. 2015.

In addition to the variability of the number of APs within bursts, BS VIP neurons also showed remarkable differences in discharge patterns in response to increasing stimulation intensities: most BS VIP neurons fired only few APs after the burst (53%, 29 of 55; Figure 1A, bottom left and right). This cease in AP firing occurred within the first 500 ms of current stimulation, leading to a prolonged phase of quiescence. These neurons showed a prominent repolarization after the burst already at rheobase stimulation. In a third of all BS VIP neurons, the burst was followed by irregular discharges of APs (29%, 16 of 55; Figure 1A, top right). The smallest group of BS VIP neurons continuously fired APs after bursting for the entire duration of current stimulation (18%, 10 of 55; Figure 1A, top left). These differences in sustained firing may result from differences in the expression of delayed rectifying K^+^ channels. VIP neurons have been shown to be equipped with such channels (Porter et al. 1998).

In response to hyperpolarizing stimuli, BS VIP neurons displayed a delayed voltage sag and slow rebound depolarizations both of variable magnitude. In all cases but one (Figure S2B) rebound depolarizations were never large enough (peak depolarization: −60.3±4.3 mV following −100 pA current stimulations) to elicit rebound spiking (see hyperpolarizing traces in Figure 1A and Figure S2). Burst firing, according to our definition, is qualitatively different from single AP firing. Therefore, we asked whether basic electrophysiological properties of BS VIP neurons (n = 55) differ substantially from those of non-BS VIP neurons (n = 214; Figure 1B, Supplementary Table 1). BS VIP neurons were more excitable in comparison to non-BS VIP neurons, with a statistically significant lower rheobase (BS: 51.6±22.0 pA; non-BS: 73.2±36.6 pA; P < 0.001). However, neither resting membrane potential (RMP; BS: −65.1±3.5 mV; non-BS: −66.3±3.6 mV; P = 0.735) nor input resistance (BS: 337.2±112.9 MΩ; non-BS: 294.2±110.9 MΩ; P = 0.09) were significantly different between BS and non-BS VIP neurons. BS VIP neurons had a significantly larger membrane time constant than non-BS VIP neurons (BS: 18.5±5.2 ms; non-BS: 15.5±5.4 ms; P < 0.001), and showed a greater voltage sag (BS: 12.1±5.5%; non-BS: 9.5±5.3%; P < 0.001). The first AP at rheobase stimulation of BS VIP neurons had a longer latency (BS: 113±41.5 ms; non-BS: 92.8±56.5; P < 0.001). The time to peak of the first AP was significantly shorter (BS: 0.367±0.054 ms; non-BS: 0.402±0.060 ms; P < 0.001) and the AP was shorter in duration than this of non-BS VIP neurons (half width of APs; BS: 0.448±0.097 ms; non-BS: 0.524±0.106 ms; P < 0.001). The time to peak of the fast AHPs following the first spike was significantly shorter in BS VIP neurons than in non-BS VIP neurons (BS: 0.497±0.127 ms; non-BS: 0.718±0.246 ms; P < 0.001). More not-significantly different membrane properties are given in Supplementary Table 1.

In conclusion, BS VIP neurons are an electrophysiologically distinct subset of layer II/III VIP neurons because they fire high-frequency bursts at any stimulus strength. This distinctness does not strictly go in parallel with differences in all basic parameters. However, since almost 50% (7 out of 16) of the basic parameters were actually statistically different, we cannot reject the possibility that BS and non-BS VIP neurons constitute different cell types also on the basis of basic membrane properties. Most of the distinct membrane properties do not intuitively relate to burst firing. Interestingly, however, the difference in rheobase (but not in RMP, input resistance, or firing threshold) indicates recruitment of additional voltage-dependent conductances. Such conductances are usually assumed to underlie bursts (Huguenard 1996).

### Location of BS VIP neurons within layer II/III

As mentioned above, the soma location of BS VIP neurons was restricted to layer II/III. We asked whether there are any differences in the soma location between BS and non-BS VIP neurons. To this end, we measured the distance of somata to the pial surface of all recorded VIP neurons (n = 269; see Material and Methods for details). We found a clear location bias of BS VIP neurons within layer II/III in that the 55 BS VIP neurons preferred upper layer II/III. The 214 non-BS VIP neurons, in contrast, were much more evenly distributed across layer II/III (Figure 1D). The median distance to the pial surface was 217 µm for BS versus 233 µm for non-BS cells (P = 0.008).

### Morphological properties of BS VIP neurons

In order to test whether BS VIP neurons are a morphologically distinct subset of layer II/III VIP neurons, we compared the morphology of 12 reconstructed BS VIP neurons to that of 8 reconstructed non-BS VIP neurons published in an earlier study (Prönneke et al. 2015). We will first describe basic morphological propertie,s followed by an analysis of the distribution of neurites throughout the barrel cortex.

BS VIP neurons were not different from non-BS VIP neurons in any of their basic morphological properties such as (i) roundness of soma (0.62±0.12), (ii) number of dendrites (2 to 5), total length (2663±716 µm), or ends of dendrites (32±11), and (iii) axonal length (8149±2392 µm) and axonal bifurcations (100±43), as well as number (2474±913) and density of axonal boutons (30±3.3 boutons per 100 µm).

These basic properties, however, do not include information about the distribution of neurites throughout the barrel cortex, which is very important for understanding the input-output relationship of VIP neurons. To describe the distribution pattern of neurites in detail, and compare BS to non-BS VIP neurons, individual neurons were aligned at the same horizontal somatic position. This method preserved the vertical distribution of their neurites (Figure 1C; S2 + S3). We then determined the average density profiles for dendritic and axonal trees for all reconstructed VIP neurons (see Material and Methods for details).

Dendrites of BS VIP neurons were found from layers I to IV with varying degrees of density. On the population level, the dendritic density of BS VIP neurons peaked in layer I and steadily decreased towards layer IV (Figure 1C, orange vertical graph, Figure S3A). Non-BS VIP neurons also spread their dendrites throughout layers I to IV. In contrast to BS VIP neurons, their dendritic density peaked in layer II/III (Figure S3B). However, this comparison was based on population data and individual VIP neurons differed in their dendritic distribution profile (Figure S5+S6). Most of the BS VIP neurons (9 of 12) had the majority of their dendrites located above their somata, in 1 neuron dendrites were distributed symmetrically with respect to the soma, and in 2 neurons dendrites were preferentially located below their somata (Figure S5). By contrast, only 1 non-BS VIP neuron had the majority of their dendrites above the soma, 3 neurons showed a symmetrical distribution, and dendrites of the remaining 4 were preferentially located below their somata (Figure S6).

The axon of BS VIP neurons extended throughout all layers of the barrel cortex. Axonal density peaked in layers II/III and Va (Figure 1C, green vertical graph). This bimodal distribution pattern is very similar to previously described layer II/III VIP neurons (Prönneke et al. 2015). Indeed, when we compared the density profiles of BS and non-BS VIP neuron, these were virtually identical (Figure S4D). This suggests that the population of BS and non-BS VIP neurons have largely overlapping output domains but does not necessarily exclude differences in target cells.

The density profiles also allowed us to analyze the horizontal spread of the neurites of BS and non-BS VIP neurons. Dendrites of BS and non-BS VIP neurons were found within ~150 µm of the soma location. Axonal trees spread further horizontally, with most of the axon found within ~250 µm. This results in the typical slender and radial appearance of BS and non-BS VIP neurons. The density of both, dendrites and axon peaked at the position of their somata and steadily decreased to both sides (Figure 1C, horizontal graphs; S2 + S3). Accordingly, the horizontal density profile of dendrites and axon was very similar when comparing BS to non-BS VIP neurons (Figure S3D + S4D).

In summary, differences in the morphology between BS and non-BS VIP neurons were at most subtle. Specifically, dendritic trees of BS VIP neurons were slightly more asymmetric and pia-oriented. Axonal trees, however, were virtually identical.

### BS VIP neurons switch their firing pattern to tonic upon depolarization

Strikingly, bursting in VIP neurons was dependent on the membrane potential. When we constantly depolarized BS VIP neurons to a membrane potential of −50 mV, their firing pattern changed to a tonic mode. In response to rheobase stimulation, all tested neurons (n = 50) fired a single action potential instead of their typical high-frequency burst (Figure 2A, top traces are responses to rheobase stimulation). Progressively increasing stimulation intensities evoked AP trains with incremental firing frequencies. In most cases the cells fired constantly (as the example shown in Figure 2A), resembling the well-known tonic mode of thalamic relay cells (Llinas and Jahnsen 1982). Note however, that some BS VIP neurons did not sustain firing at depolarized membrane potential. As will be described in detail below, even these cells fire at substantially lower frequencies compared to burst mode at RMP. Nevertheless, we will use the term tonic mode for the discharge pattern at depolarized membrane potentials.

**Figure 2:**
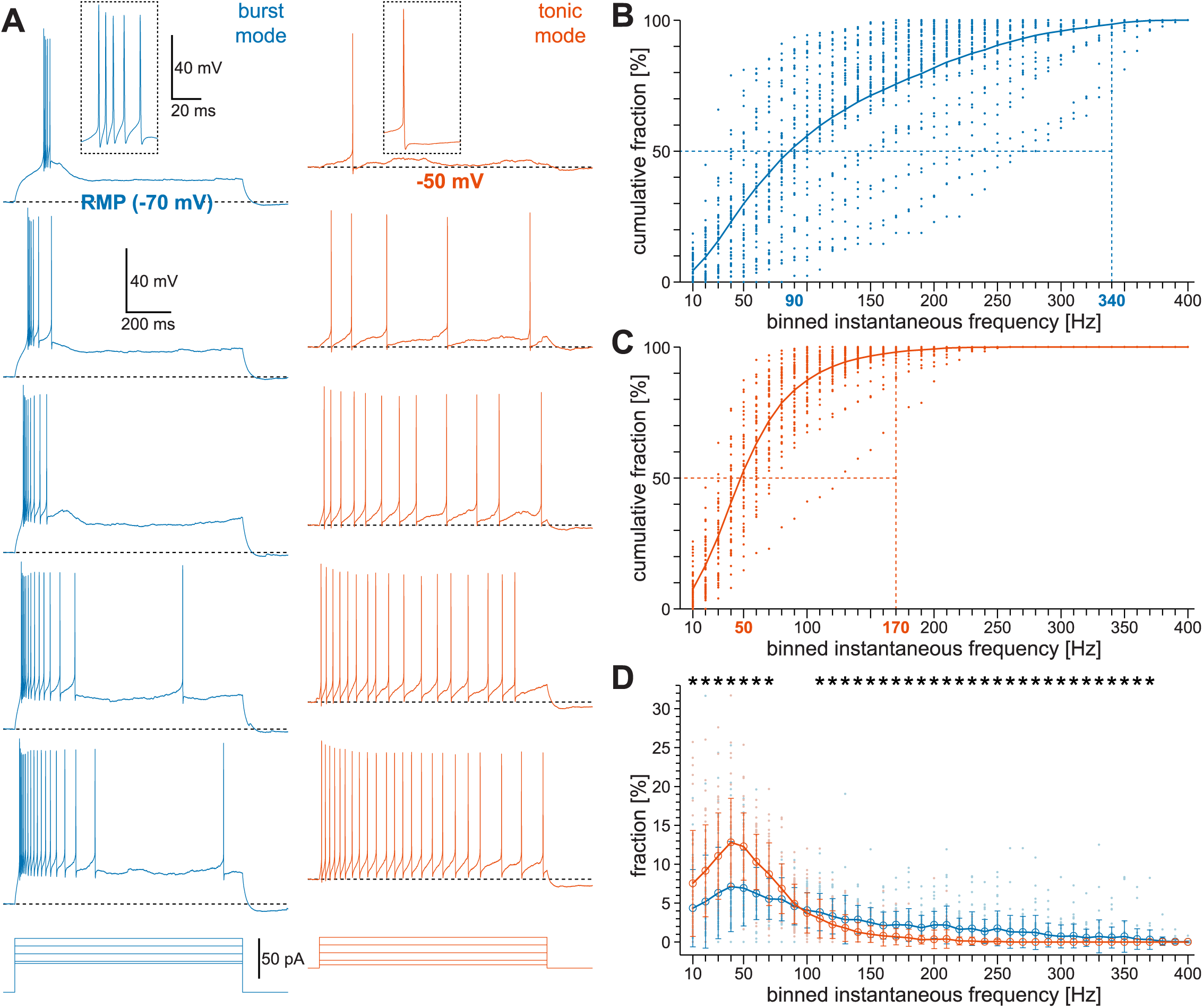
BS VIP neurons switch to a tonic firing mode upon membrane depolarization (A) Responses of a single BS VIP neuron at RMP (−65 mV; blue) and upon membrane depolarization to −50 mV (red). Top traces depict action potentials at rheobase stimulation (insets show APs with a higher temporal resolution), followed by responses to increasing stimulation strength (stimulating rectangular pulses beneath recordings). (B) Frequency spectrum of BS VIP neurons at RMP. All recorded IFFs (the reciprocal of the inter-spike interval) of individual BS VIP neurons were pooled, quantified in 10 Hz bins, and relative proportions plotted cumulatively. At RMP, 98% of all ISIs are found in a frequency range from 10 to 340 Hz (blue stippled vertical line, solid line is mean per bin, dots indicate individual data points) and the half-maximum reached at 90 Hz (stippled horizontal line). (C) Frequency spectrum of BS VIP neurons at a membrane potential of −50 mV (plot as in B). In tonic mode, 98% of all ISIs are found in a frequency range from 10 to 170 Hz and half-maximum reached at 50 Hz. (D) Comparison of the frequency spectra in burst and tonic mode. In tonic mode, the fraction of IFFs significantly inreases in the frequency range of 20 Hz to 70 Hz (red; open circles mark average per bin; error bars SD) in comparison to burst mode (blue). In the range of 110 Hz to 370 Hz, corresponding to IFFs within bursts, the fraction of IFFs significantly decreases during tonic mode.

BS VIP neurons are, therefore, able to generate two substantially different discharge patterns, especially in the frequency regime. Accordingly, we analyzed the frequency spectra by pooling the IFFs of all recorded responses of each neuron for both, burst and tonic mode separately (n = 50). We then calculated the percentile distribution of the number of IFFs in bins of 10 Hz. In burst mode (Figure 2B), the population of BS VIP neurons showed a broad range of IFFs from 10 to 340 Hz (see Materials and Methods for range definition). In tonic mode, the upper limit of the IFF range of BS VIP neurons shifted to 170 Hz (Figure 2C). On average, 50% of all IFFs were found above 90 Hz in burst mode, whereas in tonic mode 50% of all IFFs were found above 50 Hz. Thus burst mode broadens the dynamic frequency range.

A direct comparison of the frequency spectra of the population of BS VIP neurons (Figure 2D) during burst and tonic mode reveals a statistically significant difference in two separate frequency domains. Tonic mode is characterized by an increase in IFFs in the range of 10 Hz to 70 Hz and a decrease in the range of 110 Hz to 370 Hz (Figure 2D). Taking into account the IFF distribution within bursts (Figure S1) the decrease of the fraction of IFFs in the high frequency range in tonic mode might be simply explained by the absence of bursts. This argument also holds true for the increase of the fraction of IFFs in the low frequency range. The lack of post-burst spiking in some BS VIP neurons (Figure 1A) will also contribute to this.

The change in firing pattern was accompanied by slight changes in the waveform of APs and AHPs (Table S2). In tonic mode, APs had a less steep slope (burst mode: 245.7±41.2 V/s vs. tonic mode: 221.1±34.7 V/s; P = 0.02). AHPs increased in amplitude (burst mode: 10.1±2.8 mV vs. tonic mode: 12.2±3.1 mV; P = 0.008) and showed a longer time to peak (burst mode: 0.48±0.098 ms vs. tonic mode: 0.64±0.13 ms; P < 0.001).

In summary, our data show that bursting in VIP neurons was only present at a more hyperpolarized membrane potential and switched to a tonic firing mode following depolarization. This change of firing patterns substantially restricts the dynamic firing frequency range in tonic mode.

### Which mechanism underlies burst firing and burst-tonic switching in BS VIP neurons?

The characteristics of firing patterns in BS VIP neurons described so far closely resemble those of thalamic relay neurons (Llinas and Jahnsen 1982). Accordingly, we tested whether T-type calcium channels also mediate this firing behavior in BS VIP neurons by means of pharmacologically blocking these channels. In a first series of experiments (n = 5; Figure 3A), blockade of APs (0.5 µM TTX) uncovered an initial, transient depolarizing envelope (blue trace) in response to sufficiently strong current stimuli in BS VIP neurons at RMP. Additional application of the potent T-type calcium channel antagonist TTA-P2 (20 µM; (Choe et al. 2011)) blocked this transient depolarization (green trace). As summarized in Figure 3B, this holds true for all cells tested. Figure 3B also indicates a substantial variability in the magnitude of t current-induced membrane potential responses. To directly show the effect of T-type calcium channel blockade on burst firing, we applied TTA-P2 separately (n = 10) and evoked suprathreshold responses at RMP (Figure S7A). BS VIP neurons indeed switched from burst firing under control conditions to single spike firing during application of TTA-P2. In addition, blockade of t currents at RMP changed the firing frequency spectra in a similar way as depolarization of the membrane potential under control condition (compare Figure 2B-D to Figure S7B-D). Interaction of t and h currents generates rebound spiking in thalamic relay cells (Llinas and Jahnsen 1982; McCormick and Pape 1990). Although we found indications for the presence of h currents in BS VIP neurons, e.g. voltage sag and rebound depolarization in response to hyperpolarizing stimuli, we very rarely observed rebound spiking at RMP. To clarify whether BS VIP neurons actually possess HCN channels, we applied strong negative current pulses (−100 pA, 1 s) under control conditions and during application of the HCN channel antagonist ZD7288 (BoSmith et al. 1993) both at RMP. As shown in Figure 3C+D, both, the voltage sag and the rebound depolarization apparent under control condition (blue trace), fully disappear when HCN channels are blocked (purple trace). Together, these experiments show that BS VIP neurons are equipped with both T-type calcium channels and HCN channels, the former being responsible for burst firing at RMP.

**Figure 3:**
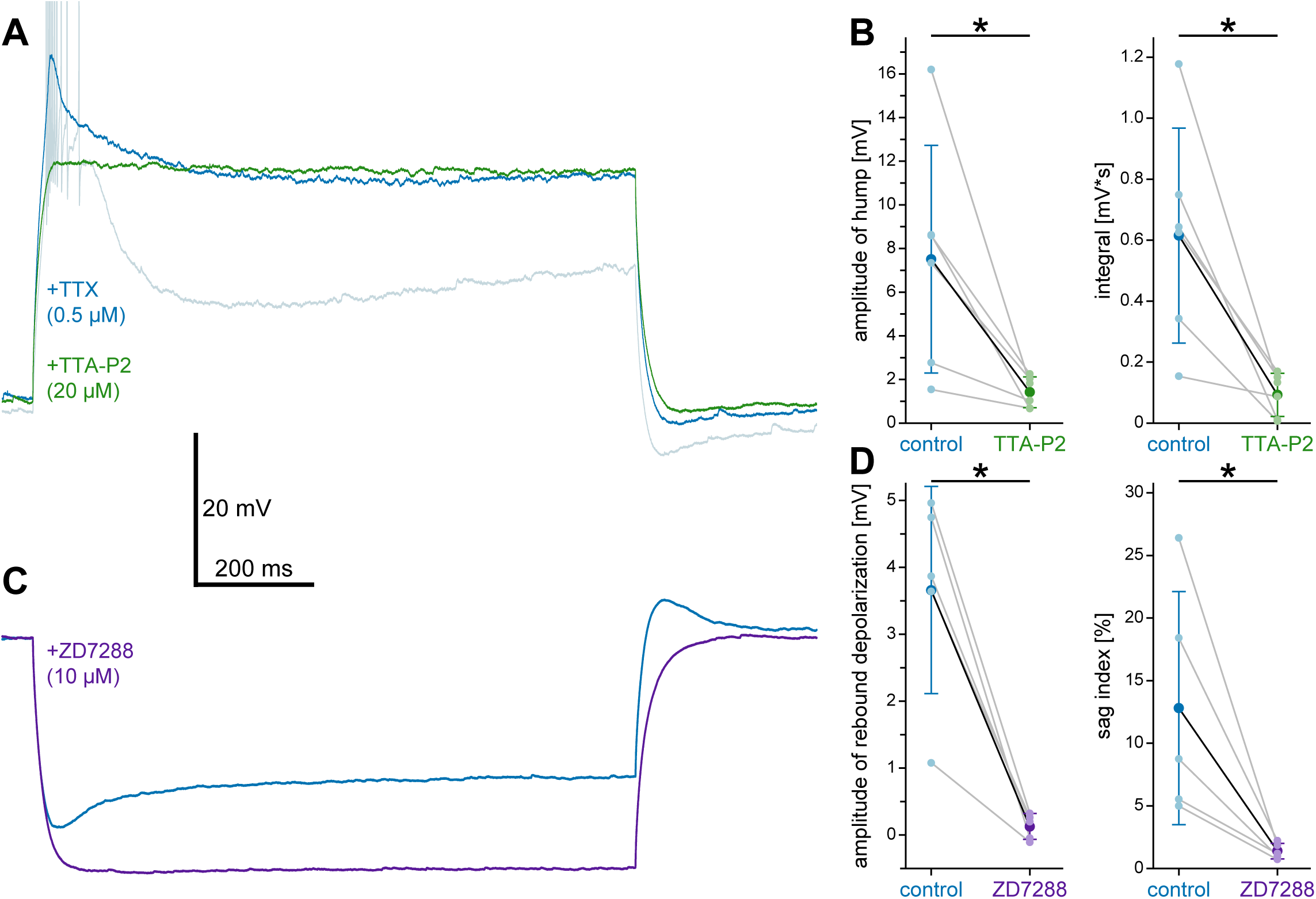
BS VIP neurons show t and h currents (A) Responses of a single BS VIP neuron at RMP (−70 mV) to depolarizing stimuli under control conditions (light blue trace, spikes are truncated), bath application of TTX (blue trace), and TTX and TTA-P2, a T-type calcium channel antagonist (green trace). (B) Quantification of the effect of TTA-P2 on amplitude (left plot, n = 5, large circles and black line indicates average, error bars SD, small circles and grey lines individual data points) and integral (right plot, n = 5) of depolarizing humps isolated by bath-applied TTX (control, blue data points). Bath application of TTA-P2 (green data points) abolished depolarizing humps as indicated by a significant reduction of amplitude and integral. (C) Responses of a single BS VIP neuron at RMP (−65 mV) to hyperpolarizing stimuli (−100 pA) under control conditions (blue trace) and during bath application of ZD7288, a HCN channel antagonist (purple trace). (D) Quantification of the effect of ZD7288 on the amplitude of the rebound depolarization following stimulation (left plot n = 5, data shown as in B) and the voltage sag (shown as sag index, right plot, n = 5). Bath application of ZD7288 abolished rebound depolarizations and voltage sags as indicated by a significant reduction (purple data points) for both events.

### Cholinergic and serotonergic modulation of VIP neurons

So far, we have characterized the burst-tonic switch of VIP neurons by depolarizing current injections. Physiologically, however, membrane potential depolarizations depend on other mechanisms, for example barrages of excitatory synaptic input or neuromodulator release. Since ACh and 5HT, two prominent neuromodulatory systems, are known to depolarize VIP neurons in primary somatosensory cortex (Porter et al. 1999; Ferezou et al. 2002) we hypothesized that their action might cause the burst-tonic switch. If so, one might ask whether the required strong depolarization is specific for BS VIP neurons, or whether all layer II/III VIP neurons are affected similarly. In a first set of experiments we bath-applied ACh (40 µM; n = 18) and 5HT (5 µM; n = 27), while monitoring the membrane potential of VIP neurons. Long-lasting bath application also enabled us to test the effects on firing patterns, i.e. the burst-tonic switch as well as the dynamic frequency range.

Both neuromodulators induced a statistically significant steady-state depolarization (ACh: P < 0.001; 5HT: P < 0.001; Figure 4A + B) in all tested VIP neurons without a significant difference in magnitude between ACh and 5HT (Figure 4A + B; ACh: 8.6±3.7 mV vs. 5HT: 7.2±3.0 mV; P = 0.19). We also monitored the recovery from the drug-induced effects during a prolonged wash-out phase. VIP neurons readily repolarized to their original membrane potential during recovery from ACh within ten minutes (Figure 4B; control: −62.8±4 mV; ACh: −54.3±5.1 mV vs. recovery: −59.7±5 mV; P = 0.003). After application of 5HT, on average, the depolarization persisted in VIP neurons for more than 30 minutes (control: −62.5±4.9 mV; during 5HT: −55.3±5.1 mV vs. recovery: −54.7±5.6 mV). None of the neurons fully recovered after 5HT application, 9 of 27 partially recovered, the remaining 18 neurons either maintained the depolarized level or even depolarized further. In addition to the long-lasting steady-state depolarizations during bath application of ACh and 5HT, we observed a fast and transient depolarization in a subset of VIP neurons during application of 5HT (Figure 4E + G) but not ACh (Figure 4C). This transient depolarization was always larger in magnitude than the steady-state depolarization and evoked spiking in most of the cells. We hypothesized that the transient responses were mediated by ionotropic 5HT_3a_R because these are fast and desensitizing (Derkach et al. 1989; Anwyl 1990; Jackson and Yakel 1995) in line with the classification of VIP neurons as 5HT_3a_R-expressing interneurons (Rudy et al. 2011). Interestingly, the persistent steady-state depolarizations outlasting the application period of 5HT point to the presence of metabotropic 5HTR (Anwyl 1990), which have not been described for cortical VIP neurons as of yet.

**Figure 4:**
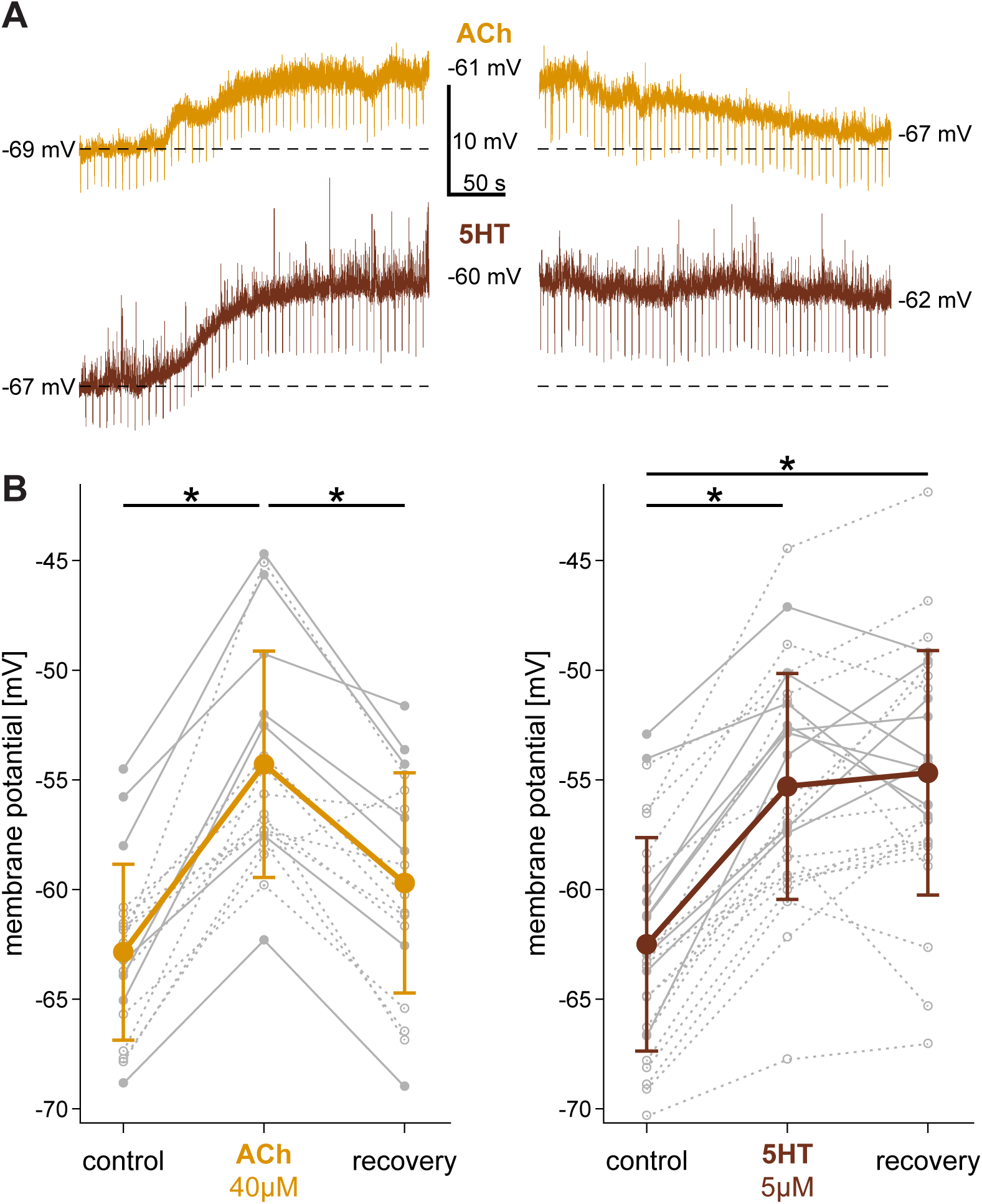
ACh and 5HT efficiently depolarize layer II/III VIP neurons (A) Example traces of two VIP neurons during bath application of ACh (40 µM; top) and 5HT (5 µM; bottom). Applications are shown on the left, recovery on the right. During recordings input resistance was monitored by applying hyperpolarizing currents (−10 pA, 200 ms duration) every 6 s. RMP is indicated to the left and a dashed line. Numbers on the right of each trace indicate the steady state membrane potentials after ten minutes of application and after ten (ACh) or thirty (5HT) minutes of recovery. (B) Summary of ACh (left; n = 23) and 5HT (right; n = 27) -induced depolarizations and recovery. Population averages±SD are shown in color, individual neurons in grey (BS VIP neurons as solid lines, non-BS as stippled). As indicated by the asterisks, ACh and 5HT significantly, and in the case of ACh reversibly, depolarized layer II/III VIP neurons.

To identify the receptors mediating the effects of ACh and 5HT, we changed our experimental protocol. VIP neurons were recorded in voltage clamp and ACh and 5HT were applied by focal pressure application. This allowed to control the stimulus duration and adding specific antagonists for receptor subtypes to the bath, while monitoring evoked currents. Short focal application (50 to 100 ms) of ACh (100 µM) reliably evoked inward currents (n = 29), which were blocked by bath-applied mecamylamine (100 µM; n = 8), an antagonist of nicotinic AChR (Figure 5A; Figure S8A). Thus, ACh acts on VIP neurons via nicotinic AChR, confirming earlier studies (Porter et al. 1999; Arroyo et al. 2012; Fu et al. 2014). Short focal application of 5HT (200 µM) evoked an influx of currents in only a subset of tested neurons (18 of 35). In some of the VIP neurons responsive to the latter approach, we combined short focal 5HT applications with bath application of tropisetron, a 5HT_3a_R antagonist (n = 5). Under these conditions, the inward current was reliably abolished (Figure 5B; Figure S8B). This finding confirmed our hypothesis that 5HT_3a_R are present in a subset of VIP neurons. Additionally, it also showed that an application period of only 100 ms does not evoke long-lasting changes in currents putatively underlying the observed steady-state depolarization.

**Figure 5:**
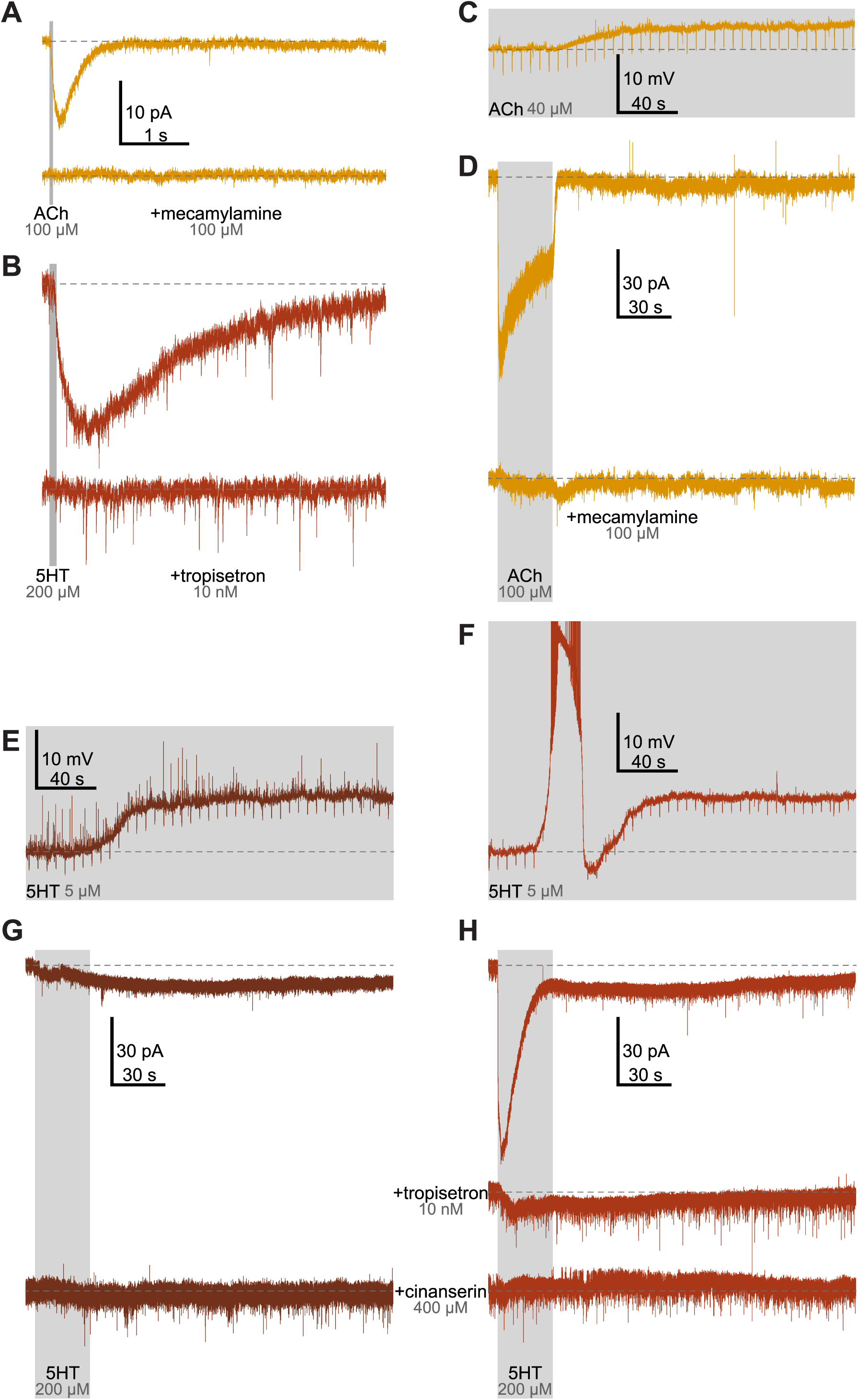
Neuromodulation by ACh and 5HT was mediated by distinct receptor subtypes The figure shows responses of layer II/III VIP neurons to focal pressure and bath application of ACh and 5HT recorded in voltage- and current-clamp. Application periods are indicated by grey rectangles, baseline holding currents or membrane potentials are always indicated by dashed lines. (A) Response of a layer II/III VIP neuron to a 50 ms-lasting focal pressure application of ACh (100 µM) recorded in voltage clamp. ACh evokes a short current influx (top trace) which is fully blocked by bath-applied mecamylamine (100 µM; bottom trace). (B) A 100 ms pressure application of 5HT (200 µM) also evokes a transient current influx (top trace). Here, bath-application of tropisetron (10 nM) fully blocks inward currents. (C) Bath-applied ACh (40 µM) depolarized a VIP neuron. The response was recorded in current clamp mode with hyperpolarizing current applications (−10 pA; 200 ms duration) every 6 seconds to monitor the input resistance. (D) Voltage clamp recording of the response of an individual VIP neurons to a 30 s focal application of ACh (100 µM; top trace). ACh evoked a strong inward current closely locked to the duration of the pressure pulse. Again, bath-applied mecamylamine (100 µM) abolished the inward current (bottom trace). (E + F) Responses of two representative layer II/III VIP neurons to bath-application of 5HT (5 µM; for details see C). While the neuron in E showed a monophasic depolarization, the one in F showed a more complex response. The latter consisted of a transient, fast-rising depolarization which was often large enough to evoke spikes (for clarity, spikes are truncated) and a persistent steady-state depolarization of lower amplitude. (G + H) Voltage-clamp recordings of the responses to focal application of 5HT (30 s; 200 µM) in two different VIP neurons corresponding to the two types shown in E and F. (G) 5HT evoked an apparent, persistent inward current with late onset (top trace) which was abolished by bath-application of cinanserin (400 µM; bottom trace). (H) In this neuron, 5HT-application evoked a strong and transient but also a persistent inward current of lower amplitude (upper trace). Bath-application of tropisetron (10 nM) abolished the transient inward current component, but not the persistent one (middle trace). The persistent inward current was blocked by bath-application of cinanserin (400 µM; lower trace).

Thus, we adjusted our focal application protocol by prolonging application time to 30 s. Under these conditions, we again observed the fast inward current described above only in a subset of VIP neurons (n = 20 of 34; Figure 5H). The decrease of the initial strong current component during application of 5HT underlines the desensitization of the respective receptor. In contrast to the current influx observed in response to short focal application, here we detected an additional inward current of low magnitude but prolonged duration (Figure 5H, upper trace). The remaining subset of VIP neuron responded to prolonged focal application only with the long-lasting current component (n = 14; Figure 5F, upper trace). Again, co-application of 5HT and tropisetron (10nM) blocked the initial fast inward current, however, it had no apparent effect on the small long-lasting inward current (n = 6; Figure 5H, middle trace; Figure S8E). If tropisetron has no apparent effect on the long-lasting component, it must be mediated by one of the metabotropic receptors. Long-lasting effects of 5HT, both depolarizations recorded in current-clamp as well as inward currents recorded in voltage-clamp, have been shown to be mediated by 5HT_2_R in many brain regions including cerebral cortex (Davies et al. 1987; Sheldon and Aghajanian 1990; Araneda and Andrade 1991; Hsiao et al. 1997; Newberry et al. 1999). Therefore, we decided to co-apply 5HT and cinanserin, a broad 5HT_2_R antagonist. As shown in Figure 5F+H (bottom traces), cinanserin (400 µM) reliably blocked the long-lasting inward currents induced by 5HT application (n = 9; Figure S8D). It has been shown that 5HT acting via the 5HT_2_R reduces the resting K^+^ leak conductance (Hsiao et al. 1997). Therefore, the apparent inward current blocked by cinanserin likely reflects a reduction of the persistent K^+^ outward current. These results are fully in line with the 5HT-evoked responses observed in current-clamp recordings described above.

When combining both datasets, only 48% (36 of 75) of VIP neurons showed 5HT_3a_R-mediated responses (Figure S9A) but all of them were affected via 5HT_2_R. Since the main focus of the current study is on BS VIP neurons, we asked whether we have the same proportional expression of the different 5HTR subtypes. We found that 69% of BS VIP neurons (16 of 23) showed 5HT_3a_R-mediated responses, while we observed this in only 38.5% of non-BS VIP neurons (20 of 52). Statistical analysis revealed a significant link between 5HT3aR-mediated responses and burst firing (Chi-square test; P = 0.025; Figure S9A). As mentioned before, however, BS VIP neurons were found more often in upper layer II/III. If VIP neurons expressing 5HT_3a_R would have a similar location bias, the higher proportion of BS VIP neurons with 5HT_3a_R might simply come by coincidence. As shown in Figure S9B, there is no such location bias for 5HT_3a_R. Therefore, 5HT_3a_R indeed are more abundant amongst BS VIP neurons.

Apparently, long focal applications are necessary to evoke responses mediated by metabotropic 5HTR. We asked whether a similar approach could also uncover metabotropic receptor-mediated responses in the case of ACh, because it also acts on ionotropic and metabotropic receptors. As shown in Figure 5D, long focal application of ACh (100 µM) induced an immediate current influx (upper trace) which persisted throughout the entire 30 s, ruling out the contribution of the alpha-7 subunit (Quick and Lester 2002). In contrast to 5HT, the current influx ceased within seconds after ACh application (n = 9). When we co-applied mecamylamine (100 µM), the observed current influx was fully blocked (lower trace; n = 6; Figure S8C). Therefore, ACh action via metabotropic receptors can be excluded.

In conclusion, ACh and 5HT substantially depolarize VIP neurons in layer II/III of the barrel cortex. While cholinergic modulation exclusively involves nicotinic receptors, serotonergic effects are more complex. In addition to the ubiquitous 5HT_2_R-mediated response, a subset of these neurons expresses functional 5HT_3a_R. Interestingly, 5HT_3a_R are much more likely to be found in BS VIP neurons than in non-BS neurons.

### Is the depolarization induced by ACh and 5HT sufficient to trigger the burst-tonic switch in BS VIP neurons?

To answer this question, we analyzed the effect of bath-applied ACh and 5HT on the firing behavior of BS VIP neurons. According to our definition, BS VIP neurons fired a burst of APs in response to rheobase stimulation (upper insets in Figure 6A + B). Regardless of the actual magnitude of depolarization induced by either neuromodulator in individual neurons (ACh n = 8; 5HT n = 9), ACh or 5HT consistently changed the response to rheobase stimulation from bursts to single spike firing (lower insets in Figure 6A + B). In any case, the initial burst pattern was recovered (insets in Figure 5E + F) when DC was injected in order to compensate for the drug-induced depolarization. Next, we asked whether depolarizations induced by ACh or 5HT influence the entire frequency spectra of BS VIP neurons in a similar way as artificial alterations of the membrane potential by DC application does (Figure 2B - D). To this end, we again applied series of suprathreshold current pulses with increasing amplitudes under control and test conditions. As described above, we calculated IFF plots from these data. Under control conditions, BS VIP neurons typically showed the initial high frequency firing followed by occasional APs at irregular intervals or even no further APs (Figure 6A + B upper traces). Besides a substantial depolarization and the disappearance of the initial burst, application of either ACh or 5HT resulted in continuous AP firing that lasted throughout the entire stimulus period (Figure 6A + B lower traces). This purely qualitative description is reflected in the IFF plots (Figure 6C + D; see Figure S10 for corresponding cumulative plots): both, ACh and 5HT, significantly increase the fraction of IFFs of lower frequency and decrease the fraction of IFFs of higher frequency. Note the similarity to the effect of DC-induced depolarization on the frequency spectra (Figure 2D). This again suggests that depolarization per se determines the firing behavior of BS VIP neurons. If so, DC compensation of the drug-induced depolarization should not only recover the initial burst but also the pattern of subsequent firing. As shown in Figure 6E + F, this approach indeed recovered the initial burst, however, the post-burst firing pattern did not fully resemble the one observed under control conditions in individual neurons. This might indicate that ACh and 5HT do not only affect conductances responsible for steady-state membrane potential. Indeed, 5HT is known to modulate voltage-gated K^+^ channels via 5HT_2_ R (D’Adamo et al. 2013). As mentioned above, VIP neurons express delayed rectifying K^+^ channels, i.e. Kv1.1 and Kv1.2, controlling sustained firing (Porter et al. 1998). Comparing the frequency spectra from cells recorded at control conditions with those recorded in the presence of ACh or 5HT but clamped to RMP (Figure 6G + H) revealed much less significantly different frequency bins as compared to the differences between control conditions and ACh or 5HT without compensation (Figure 6C + D). However, the power of statistical tests was below the desired value (0.8). Thus, we cannot reliably exclude the possibility of subtle differences here.

**Figure 6:**
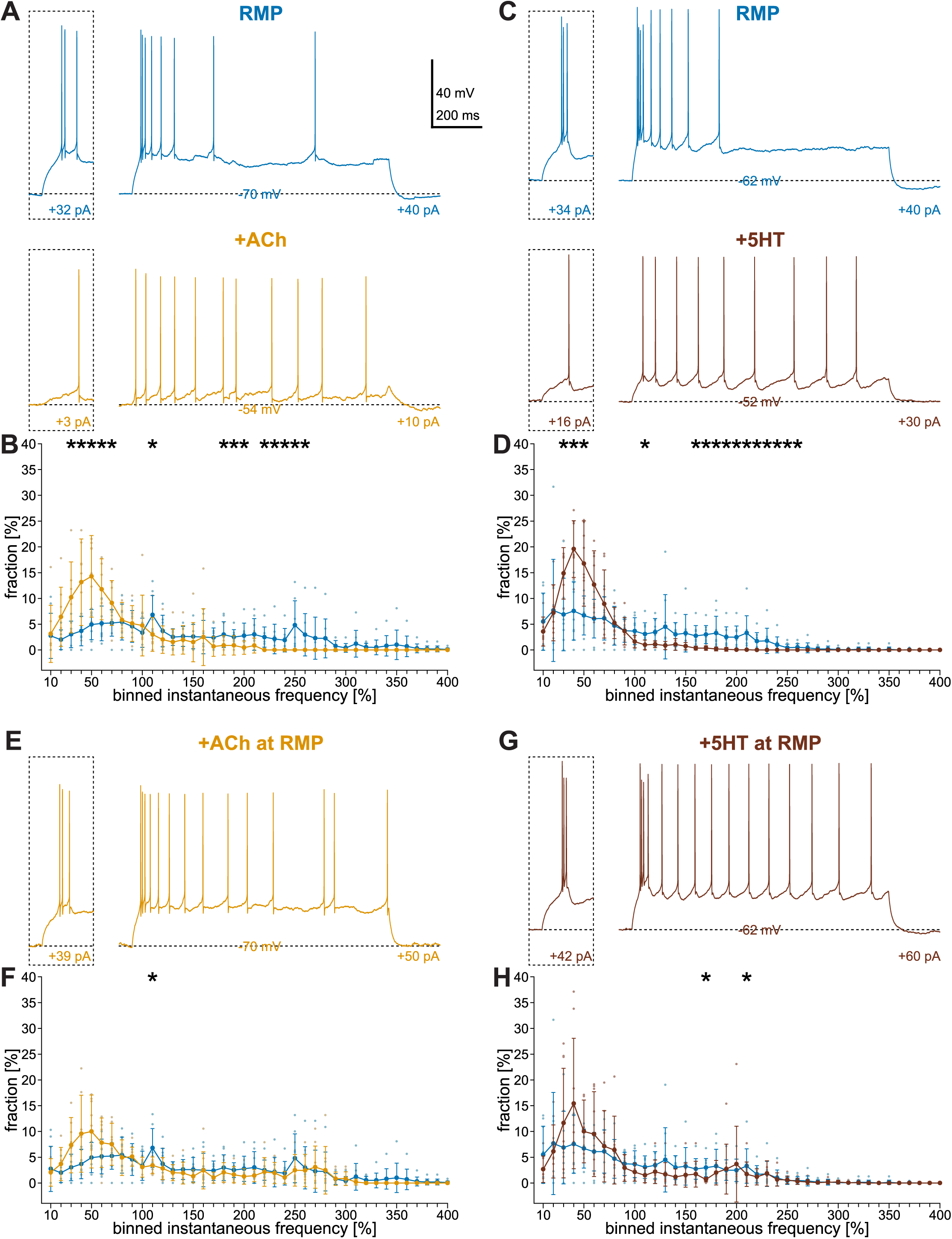
Depolarization induced by ACh and 5HT suffices to switch firing modes from burst to tonic in BS VIP neurons Effect of neuromodulation on the firing behavior of BS VIP neurons. The figure shows responses of individual neurons to suprathreshold current injections (1 s; current amplitudes are indicated below each trace) under control conditions (blue) and during bath-applications of ACh (40 µM; orange) and 5HT (5 µM; brown). The insets show the responses to rheobase stimulations, baseline membrane potentials are indicated by dashed lines and numbers. It also shows comparisons of frequency spectra. (A) The firing behavior of a BS VIP neuron at RMP (−70 mV; top traces) was changed from bursting, initial burst followed by irregular spiking, to tonic when depolarized to −54 mV by ACh (bottom traces). (B) Comparison of the frequency spectra (n = 8; color code as in A) revealed significant differences in two separate IFF ranges: ACh increased the proportion of ISIs between 30 – 70 Hz and decreased it between 180 – 260 Hz. (C) Bath-application of 5HT depolarized the example neuron from −62 mV to −52 mV. Again, this was accompanied by a switch from burst (upper traces) to tonic firing (lower traces). (D) Comparison of the frequency spectra (n = 8; color code as in C) again showed significant differences in two separate IFF ranges: 5HT increased the proportion of ISIs between 30 – 50 Hz and decreased it between 160 – 260 Hz. (E) Repolarization to the initial membrane potential of the example neuron shown in A by DC application during bath-application of ACh restored burst firing. Note however, that the cell fired more spikes after the initial burst. (F) Comparison of the frequency spectra (n = 8; color code as in A) between control condition and ACh application compensated for the depolarization showed almost none significantly IFF bins. This confirms the restoring of the burst mode. (G) DC compensation of the 5HT-induced depolarization also restored burst firing in the example cell shown in C. Again, the firing behavior after the initial burst was not fully recovered. (H) When DC was used to compensate for the depolarization induced by 5HT, the number of significantly different IFF bins was massively reduced as compared to D (n = 8; color code as in C).

## Discussion

In the present study, we describe a bimodal firing behavior in a substantial proportion of VIP neurons in layer II/III of mouse primary somatosensory cortex for the first time. These neurons switch from burst to tonic firing dependent on the membrane potential and its effect on the availability of T-type calcium currents, respectively. In contrast to this clear-cut difference in firing behavior between BS and non-BS VIP neurons, morphology and basic electrophysiological properties are strikingly similar. Furthermore, we show that both ACh and 5HT contribute to the depolarization required for such a switch. Interestingly, we discovered a so far unknown serotonergic modulation of layer II/III VIP neurons via type 2 metabotropic receptors.

### Mechanisms of the firing behavior in BS VIP neurons

Bursting in VIP neurons is characterized by two main features: (i) it occurs as an all-or-none response and (ii) its voltage-dependent switch to tonic firing. Both features can be explained by T-type calcium channels, which we show to be present in BS VIP neurons. At relatively hyperpolarized membrane potentials, T-type calcium channels are transiently activated upon depolarization and allow the generation of a burst. The lack of deinactivation of these channels at depolarized membrane potentials leads to tonic firing (Huguenard 1996). Burst firing behavior like that but also the underlying mechanism is well known for thalamic relay cell (Llinas and Jahnsen 1982; McCormick and Pape 1990). Based on a mixed action of T-type calcium channels and HCN channels, these thalamic cells show rebound burst firing enabling rhythmic activity. Although BS VIP neurons also possess HCN channels, as shown in the present study, we rarely observed rebound spiking in this population. As reviewed in (Huguenard 1996), rebound spiking based on t current strongly depends on ambient ion concentrations, deinactivation and activation thresholds. This opens several possibilities why rebound spiking occurs so rarely in our sample. If VIP neurons do not differ from thalamic relay cells in terms of surface charges, ambient ion concentrations should not be a major factor because thalamic relay cells do show rebound spiking under the conditions used here (Guy et al. 2016). T-type calcium channels require sufficient hyperpolarization to be released from inactivation. As shown in Figure S2, the hyperpolarizations evoked in the present study met this requirement both in magnitude and duration (Coulter et al. 1989). Therefore, lack of deinactivation is probably not the reason for the sparseness of rebound spiking. The threshold for activation, finally, differs substantially between cell types. In thalamic relay cells it is around −60 mV, whereas it is 9 mV more depolarized in thalamic reticular cells (Huguenard 1996). Such differences were shown to be due to differences in channel isoforms (Iftinca et al. 2006; Cain and Snutch 2010). The peak membrane potential level of depolarizing humps in just-subthreshold traces was on average around −61 mV (if corrected for liquid junction potential) in our sample (Figure S2). Although this seems to be close to the values reported in the literature, this approach is by far no precise estimation of the actual activation threshold. However, we believe that it is fair to compare the peak membrane potential of the depolarizing hump to the peak rebound depolarization. Since the latter is around 15 mV more hyperpolarized, it is not a surprise that rebound depolarizations will not consistently evoke rebound spikes. Although BS VIP neurons possess both T-type calcium and HCN channels, the striking difference between apparent activation threshold and rebound depolarizations does not put them into the position to generate autogenic rhythmic bursting.

### Bursting in VIP neurons as a classifier

Classification of cell types intends to define clear-cut groups of cells based on their intrinsic properties which will, in the best case, reveal functionally distinct populations. This holds true also for the distinction of subgroups within specific cell-types. This approach is traditionally based on large samples of basic parameters of individual neurons (Tyner 1975). Cell classes, however, are not an end in itself but should at best guide experimenters. In the present study, we described a single feature of a subpopulation of VIP neurons in layer II/III, namely bursting, which is not only a qualitative difference to other VIP neurons but can also be easily tested during actual experiments. Nevertheless, it is of interest to analyze and compare multiple basic properties. In a previous study (Prönneke et al. 2015), we described differences between VIP neurons in layer IV-VI and layer II/III based on basic electrophysiological and morphological properties. However, we did not attempt to further subdivide VIP cells in layer II/III because of a low sample size and a remarkable heterogeneity. Based on a larger sample size, especially for BS VIP neurons, we were now in the position to compare the standard parameters of BS VIP neurons to those of non-BS VIP neurons in layer II/III. In terms of morphology, the most striking result was the restriction of somata of BS VIP neurons to the upper two thirds of layer II/III, a finding that will also simplify further studies on BS VIP neurons. The distribution of neurites was almost identical for both groups. As mentioned above, this does not indicate shared input sources and output targets. In terms of electrophysiology, we found more differences between the two groups, however, they are not fully distinct. Moreover, it requires post-hoc analysis of a large sample to reveal putatively separate cell types, which could have been already identified based on the occurrence of burst firing or not.

Since our analysis was based on classical electrophysiological and morphological properties, it is important to also consider molecular approaches to classify VIP neurons, especially regarding transcription of the genes relevant for burst spiking. We identified T-type calcium channels as the mechanism for burst spiking in BS VIP neurons. These channels contain one of three different alpha-subunits (McRory et al. 2001), encoded by the genes CACNA1G, CACNA1H, and CACNA1I. To our knowledge, there is only one report describing single cell transcriptomics in mouse somatosensory cortex (Zeisel et al. 2015), however, none of these genes were reported for interneurons in this study. In a recent study in mouse visual cortex, the population of VIP neurons was subdivided into 5 subgroups based on mRNA expression profiles (Tasic et al. 2016). In this study, the expression of T-type calcium channel encoding genes is neither restricted to 1 of the subgroups of VIP neurons, as defined by Tasic and colleagues, nor is it a common feature of all neurons within any of these subgroups. On the other hand side, CACNA1G-I expression is also not equally distributed or at least detected in a substantial number of neurons across each of the subgroups. Thus, expression of T-type calcium channel encoding genes does not clearly match to subgroups identified by single cell transcriptomics so far.

### Mechanisms of cholinergic and serotonergic modulation

In the present study, we showed that both BS and non-BS VIP neurons in layer II/III are affected by ACh and 5HT. While ACh acts via nicotinic receptors, 5HT action is mediated by ionotropic 5HT3aR and metabotropic 5HT_2_R. Modulation of VIP neurons by ACH and 5HT has been shown before (Porter et al. 1999; Ferezou et al. 2002; Arroyo et al. 2012; Fu et al. 2014), however, without focus on putatively distinct subtypes. As we demonstrate here, BS and non-BS VIP neurons show similar responses to both neuromodulators. In addition to confirming previous studies, we found two novel aspects: (i) serotonin ubiquitously operates via metabotropic receptors, and (ii) ionotropic serotonin receptor-mediated responses are lacking in more than half of the layer II/III VIP neurons. The latter finding seems somewhat surprising since VIP neurons are considered to be a subgroup of 5HT3aR-expressing interneurons (Lee et al. 2010; Rudy et al. 2011). As long as this controversy is not resolved, 5HT3aR-expression as a marker for VIP neurons should be taken with caution.

From a functional point of view, the disparity in 5HT3aR-mediated responses is of interest because it allows a differential recruitment of VIP neurons by 5HT. This is even more striking, because we have shown in the present study that a significantly larger proportion of BS VIP neurons possess functional 5HT3aR than non-BS VIP neurons. ACh and 5HT, the latter via metabotropic receptors, modulate the membrane potential of all VIP neurons indiscriminative according to the brain state. By contrast, 5HT3aR, characterized by rapid desensitization and slow recovery from it (Derkach et al. 1989; Anwyl 1990; Jackson and Yakel 1995), might introduce a more temporally structured modulation of only a subset of VIP neurons. Noteworthy, we do not argue that the effect mediated by nicotinic AChR and 5HT_2_R on individual cells is indiscriminative. Due to the nature of metabotropic receptors and in line with our experiments, 5HT will only affect VIP cells via 5HT_2_R in vivo if the corresponding dorsal raphe neurons exhibit adequately sustained and high-frequent activity. Previous studies have indeed shown that serotonergic neurons in the dorsal raphe are highly active during waking and slow wave sleep with average firing frequencies ranging from 9 to 75 Hz (Kocsis and Vertes 1992; Kocsis et al. 2006). Since released ACh is quickly hydrolyzed (Ellman et al. 1961), also cholinergic inputs to VIP neurons acting via nicotinic AChR require high activity of basal forebrain neurons in order to generate prolonged periods of depolarization. ChAT-positive basal forebrain neurons in the awake animal show ongoing activity at instantaneous firing frequencies of more than 100 Hz on average (Lee et al. 2005; Hassani et al. 2009).

### Potential functional relevance of burst spiking in VIP neurons

The ability of a substantial proportion of layer II/III VIP cells to work in two completely distinct firing modes, burst and tonic firing, potentially dependent on brain states may allow them to have a specific functional impact on cortical circuitry distinct from all other VIP cells. Firing frequency influences communication between neurons (Lisman 1997; Owen et al. 2018). The burst in BS VIP cells consists of two to several spikes with roughly constant instantaneous frequencies regardless of input strength. By contrast, the firing frequency in tonic mode varies much more with input strength. Therefore, BS VIP cells have a more stable output signal in burst mode and a more modifiable in tonic mode. A second aspect of the differences in firing frequencies between the two modes directly concerns synaptic transmission. High-frequency bursts are much more likely to result in presynaptic facilitation or depression (Caputi et al. 2009; Walker et al. 2016). Moreover, they allow more effective temporal summation in the postsynaptic neuron (Karnani et al. 2016).

Since BS VIP neurons operate in two different modes, it is of functional importance to reveal whether these modes are the signatures of different behavioral states or might occur in concert. In the classical case of burst-tonic switching neurons, thalamic relay cells, it was initially assumed that bursting occurs during sleep and tonic firing in the awake state (McCormick and Huguenard 1992) but a number of studies have meanwhile shown the co-occurrence of bursts and tonic firing in different awake behavioral states (Fanselow et al. 2001; Urbain et al. 2015). Simply because we report a burst-tonic switch in layer II/III VIP neurons for the first time, corresponding in vivo studies do not exist as of yet. If a direct confirmation of burst firing in the awake animal is lacking, is there any evidence for phases of hyperpolarization, the prerequisite for bursting, in layer II/III VIP neurons in vivo? Gentet and colleagues (Gentet et al. 2012) recently provided intracellular recordings from excitatory and various inhibitory neurons in layer II/III of primary somatosensory cortex of awake mice. Most of these neurons, including VIP cells, indeed show membrane potential fluctuations with prolonged periods of hyperpolarization well below −60 mV. In case that BS VIP neurons are not an exception, burst and tonic firing might both occur in the awake animal determined by these periodic membrane potential fluctuations. Clustered activity of cholinergic basal forebrain (Lee et al. 2005; Hassani et al. 2009) and serotonergic dorsal raphe neurons (Kocsis and Vertes 1992; Kocsis et al. 2006) as well as the periodic activity of excitatory neurons could possibly drive such membrane potential fluctuations in BS VIP neurons in a coordinated and behavior-dependent fashion.

## Supporting information

## Acknowledgements

We thank Patricia Sprysch for excellent technical assistance and Anette Mertens, Adrian Villalobos, Megha Patwa, and Nicolas Zdun reconstructing morphology. This study was supported by a grant from the Deutsche Forschungsgemeinschaft via the CRC 889 (Cellular mechanisms of sensory processing) subproject 07 (to JFS).

