## Supplementary material for "Neuromodulation leads to a burst-tonic switch in a subset of VIP neurons in mouse primary somatosensory (barrel) cortex"

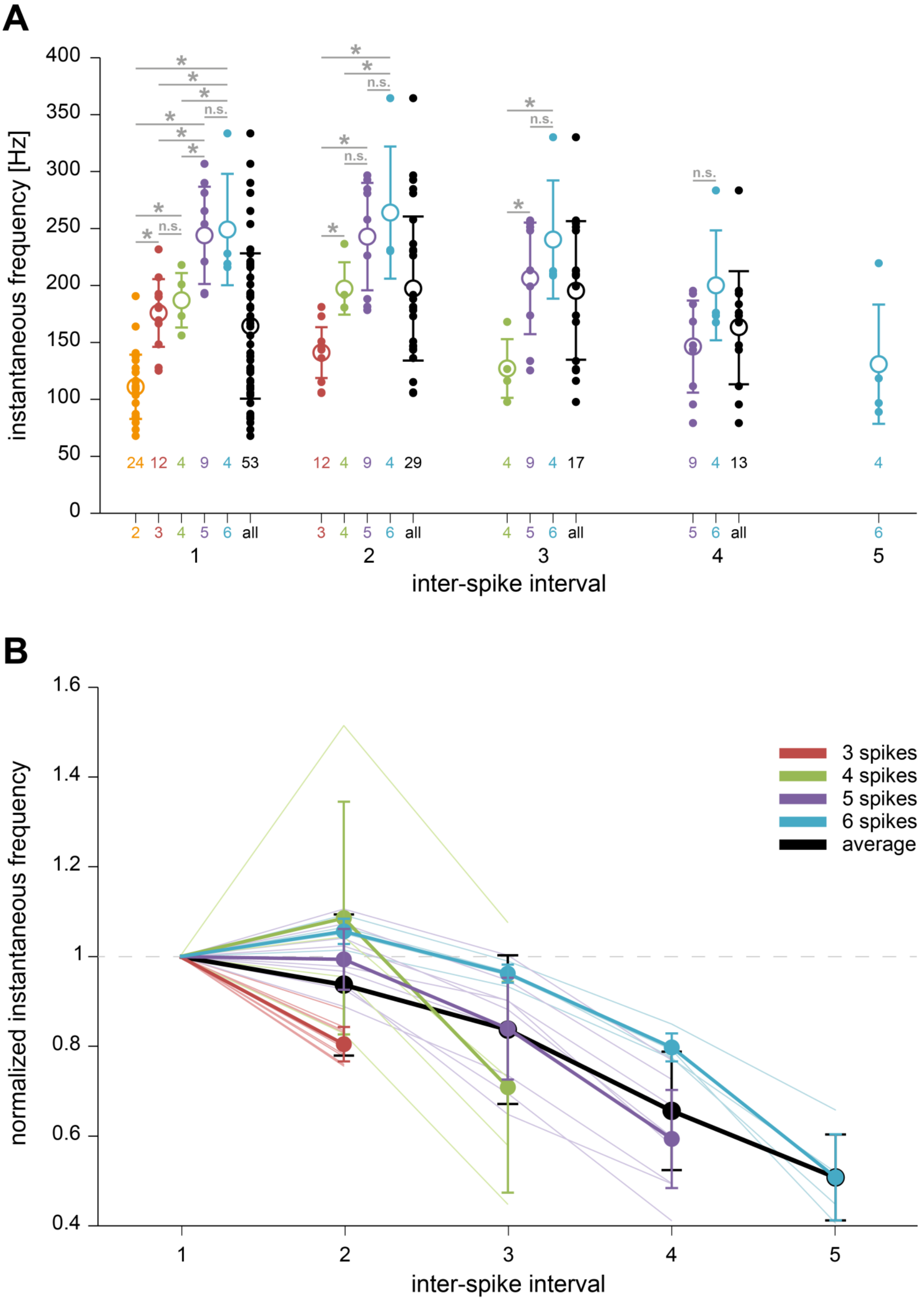
A
instantaneous frequency [Hz]
400
350
300
250
200
150
100
50
0
*
n.s.
24 12 4 9 4 53
12 4 9 4 29
4 9 4 17
9 4 13
4
2 3 4 5 6 all
3 4 5 6 all
4 5 6 all
5 6 all
6
1
2
3
4
5
inter-spike interval
B
normalized instantaneous frequency
1.6
1.4
1.2
1
0.8
0.6
0.4
3 spikes
4 spikes
5 spikes
6 spikes
average
1
2
3
4
5
inter-spike interval

Supplementary Figure 1: Detailed analysis of bursts of BS VIP neurons at rheobase stimulation

(A) Summary plot of instantaneous firing frequencies (IFF) across all inter-spike intervals (ISI) in bursts and neurons at rheobase stimulations. Filled circles indicate ISIs of individual cells, open circles and error bars indicate average±SD. For each ISI, black symbols show all individual IFFs and the average across them, colored symbols show IFFs grouped by the number of spikes per burst (see also colored label of the x-axis). The number of cells per group is given beneath data points. Asterisks indicate significant differences (one way ANOVA; n.s. = non-significant). Note that IFFs within each ISI depend on the number of spikes per burst: the more spikes per burst, the higher the IFF. See table below for details. (B) Changes in IFFs during bursts. Data shown in A were normalized with respect to the first ISI and plotted vs. ISI number. Grand average±SD is shown in black, averages±SD and individual cells are shown in thick and thin lines, respectively, color-coded according to the number of spikes per burst. Overall, IFFs decrease throughout the burst, however, bursts consisting of more than 3 spikes, on average, accelerated or did not change in the second ISI.

| Spikes | (n) | ISI 1 [Hz] | ISI 2 [Hz] | ISI 3 [Hz] | ISI 4 [Hz] | ISI 5 [Hz] |
|---|---|---|---|---|---|---|
| 2 | (24) | 110.9±28.8 | - | - | - | - |
| 3 | (12) | 175.8±31 | 141.1±23.3 | - | - | - |
| 4 | (4) | 186.9±27.7 | 197.4±26.6 | 127.1±29.7 | - | - |
| 5 | (9) | 244.0±43.3 | 242.9±50.0 | 206.2±52.0 | 146.3±42.9 | - |
| 6 | (4) | 249.1±56.5 | 264.0±66.9 | 240.3±59.9 | 200.1±55.6 | 130.9±60.4 |
| all | (53) | 164.4±64.4 | 197.4±64.4 | 195.6±62.7 | 162.9±51.7 | - |

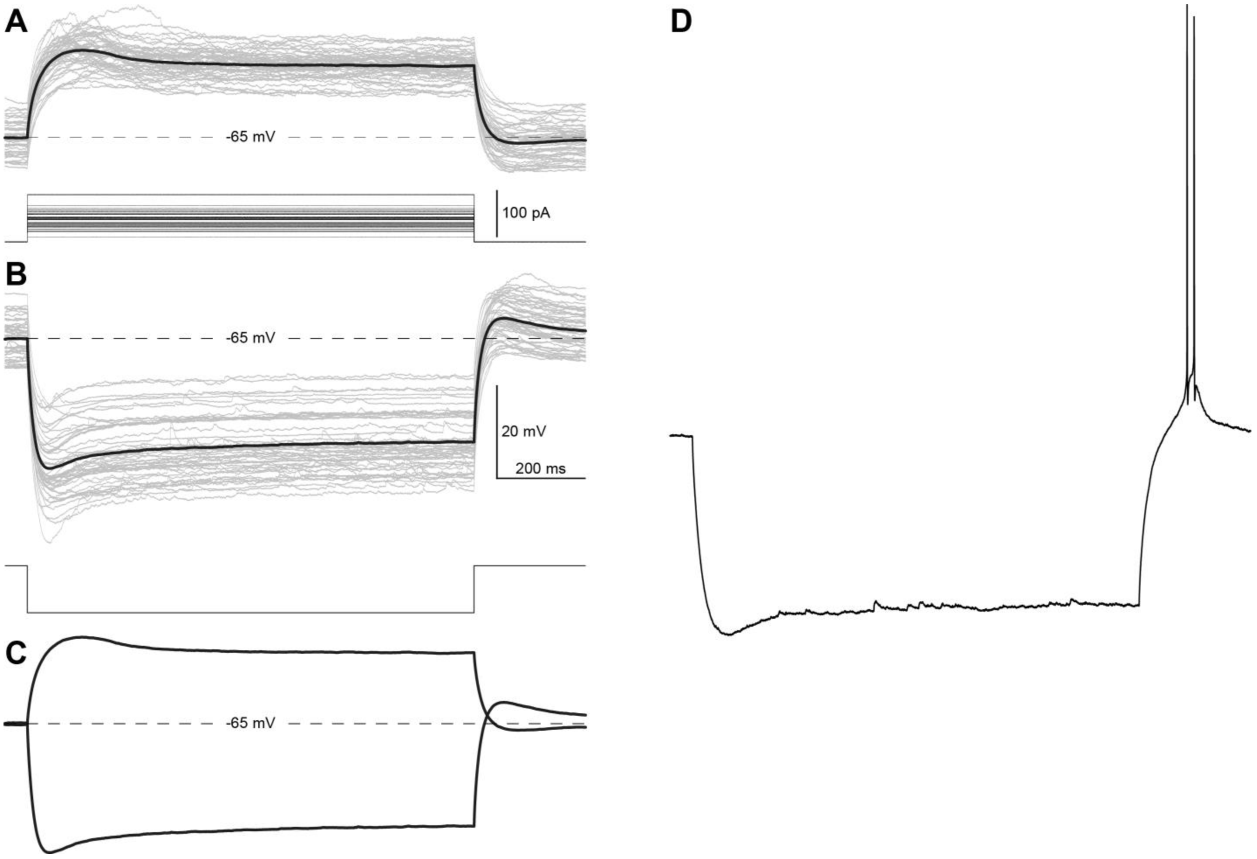


Supplementary Figure 2: Depolarizing humps and rebound depolarizations of BS VIP neurons

(A) Superposition of individual membrane potential responses (n = 49; grey traces) to just-subthreshold stimulations (1 s; 1 pA below rheobase) and average of all responses (black trace). Individual BS VIP neurons (n = 49) showed depolarizing humps of different magnitude. On average, depolarizing humps peaked at -45.3±3.5 mV (corrected for liquid junction potential: -61.3±3.5 mV). (B) Superposition of individual traces (n = 50; grey) and the average (black) in response to -100 pA current stimulation (1 s). BS VIP neurons showed moderate voltage sags and small rebound depolarizations. On average, the hyperpolarizations peaked at -93.4±6.7 mV (corrected for liquid junction potential: -109.4±6.7 mV) and reached a steady state at -87.3±6.2 mV (corrected for liquid junction potential: -103.2±6.2 mV). Rebound depolarizations following current stimulation peaked at -60.3±4.3 mV (corrected for liquid junction potential: -76.3±4.3 mV). (C) Overlay of the averages from A and B illustrating more clearly the differences between the peaks of depolarizing humps and rebound depolarizations (15±5.3 mV). (D) The only existing example of rebound spiking in our dataset.

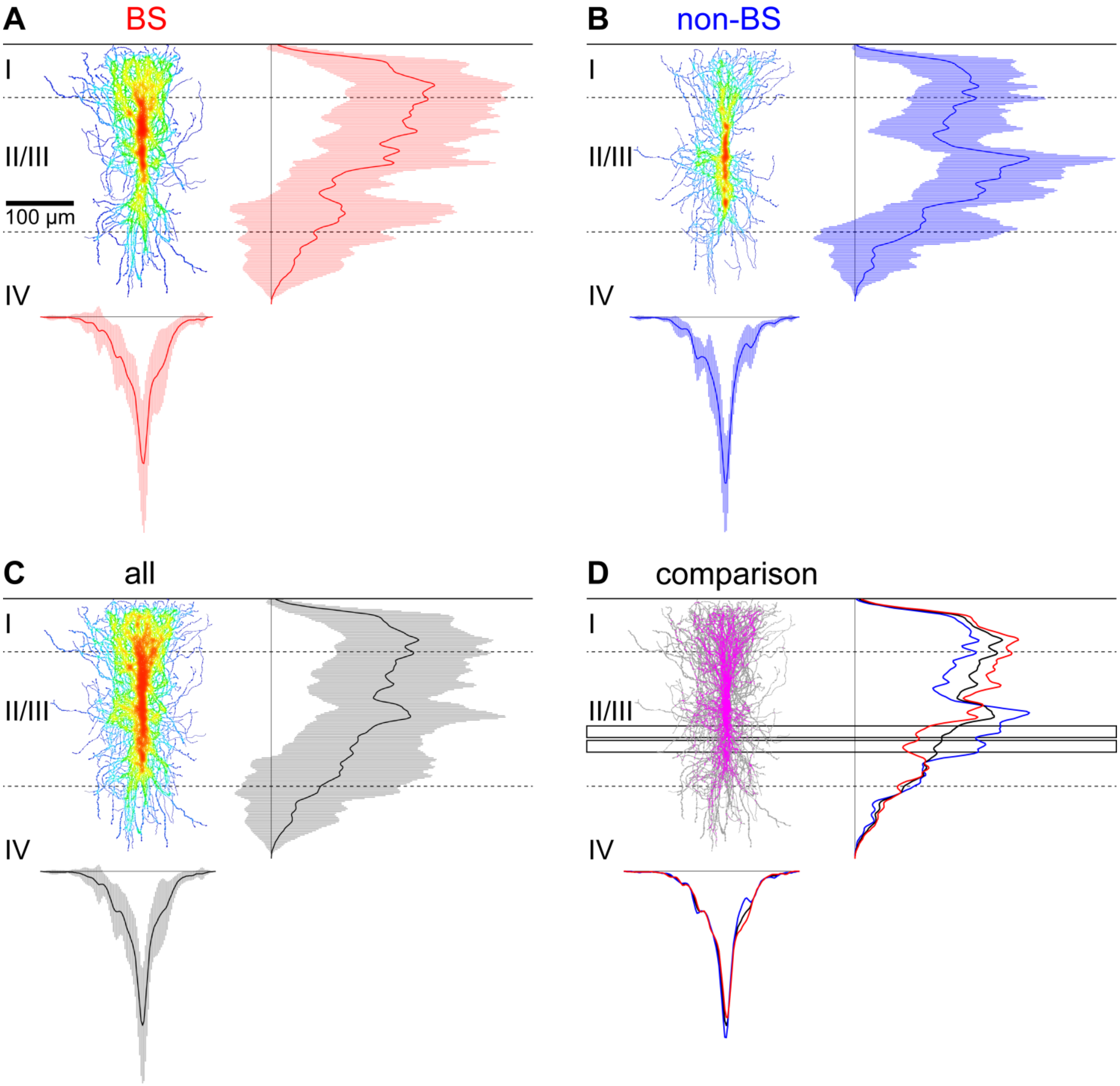

A
BS
B
non-BS
C
all
D
comparison
I
II/III
IV
100 µm

Supplementary Figure 3: Comparison of dendrites between BS and non-BS VIP neurons

(A) Superposition of dendritic trees of 12 individual BS VIP neurons (shown in Figure S5). Density of dendrites is shown as a heat-map ranging from least dense (cold colors) to most dense (warm colors). Vertical (right) and horizontal (bottom) distribution profiles plotted as averages (red traces) and error bars (SD; light red areas, Roman numerals indicate layers, dashed lines layer borders). (B) Superposition of 8 individual non-BS VIP neurons (shown in Figure S6) visualized as in A (but distribution profiles in blue). (C) Superposition of all 20 neurons shown in A and B with distribution profiles in black. (D) Direct comparison of the distribution of dendrites between BS and non-BS VIP neurons. Binary images used for the heat-maps in A and B were converted to greyscale and multiplied. Overlapping areas were pseudo-colored magenta. The averages of the vertical distribution profiles from A, B, and C are shown to the right, corresponding averages of the horizontal distributions are shown below. Areas of significantly higher dendritic density of non-BS VIP neurons were found only in the vertical distribution and are indicated by open rectangles.

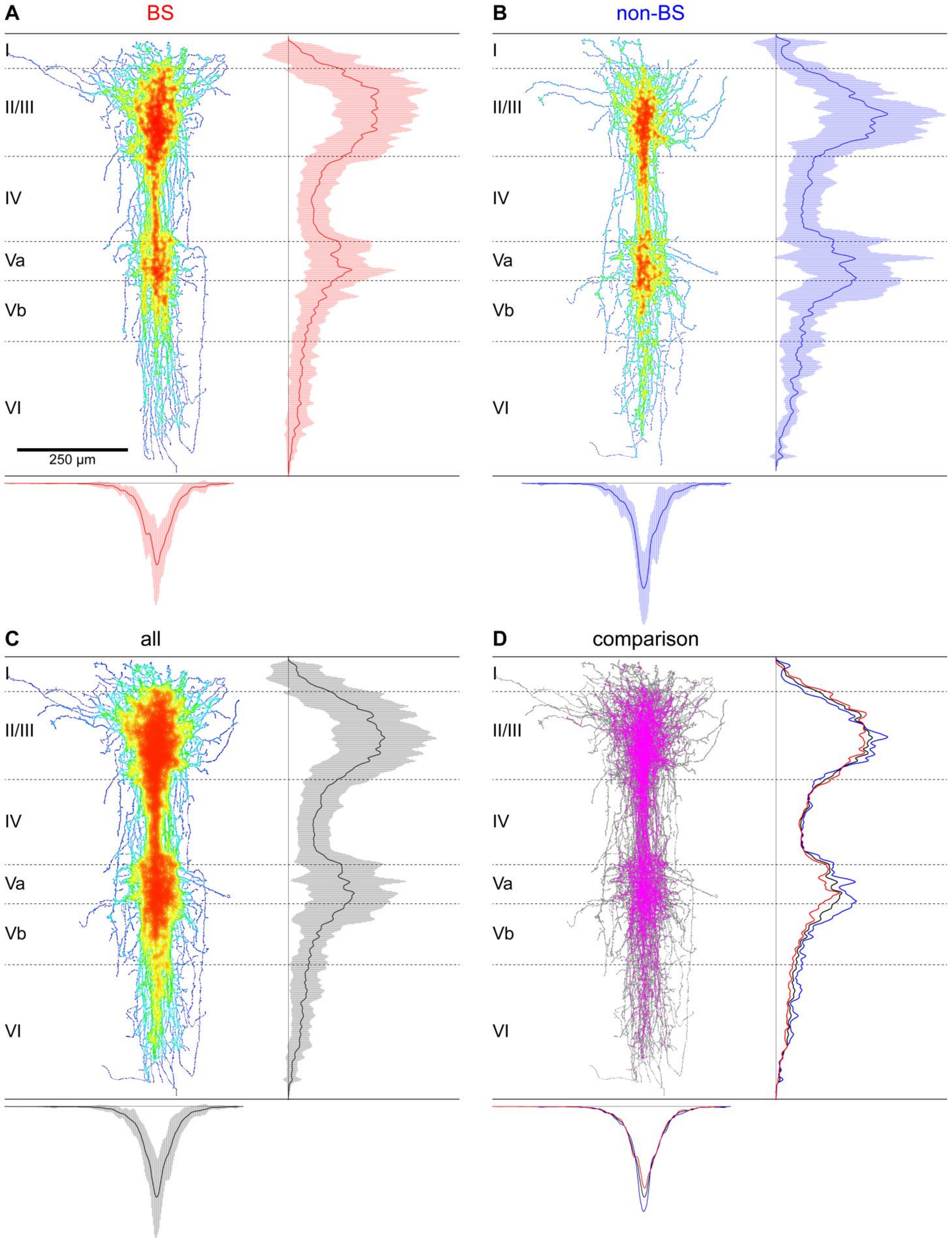

A
BS
B
non-BS
C
all
D
comparison
I
II/III
IV
Va
Vb
VI
250 µm

Supplementary Figure 4: Comparison of axon between BS and non-BS VIP neurons

(A) Superposition of axonal trees of 12 individual BS VIP neurons (shown in Figure S5). Density of axon is shown as a heat-map ranging from least dense (cold colors) to most dense (warm colors). Vertical (right) and horizontal (bottom) distribution profiles plotted as averages (red traces) and error bars (SD; light red areas, Roman numerals indicate layers, dashed lines layer borders). (B) Superposition of 8 individual non-BS VIP neurons (shown in Figure S6) visualized as in A (but distribution profiles in blue). (C) Superposition of all 20 neurons shown in A and B with distribution profiles in black. (D) Direct comparison of the distribution of axon between BS and non-BS VIP neurons. Binary images used for the heat-maps in A and B were converted to greyscale and multiplied. Overlapping areas were pseudo-colored magenta. The averages of the vertical distribution profiles from A, B, and C are shown to the right, corresponding averages of the horizontal distributions are shown below. Note that axonal trees were virtually identical between BS and non-BS VIP neurons in their vertical and horizontal distribution profile.

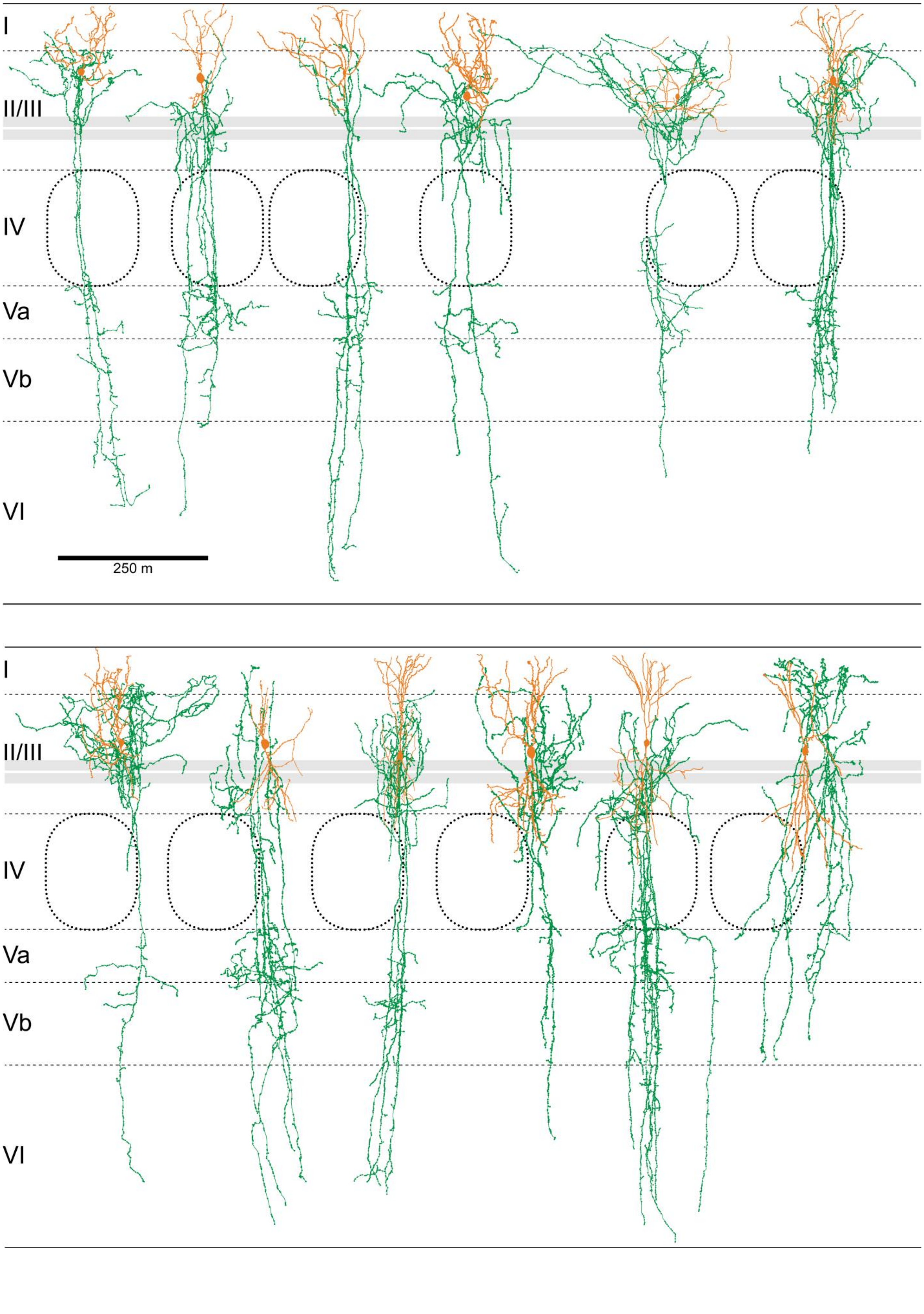
I
II/III
IV
Va
Vb
VI
250 m
I
II/III
IV
Va
Vb
VI

Supplementary Figure 5: Reconstructions of the morphology of 12 individual BS VIP neurons

The 12 individual BS VIP neurons are shown with somata and dendrites in orange and axon in green with their corresponding home barrel in layer IV. Grey rectangles indicate the area in which their dendritic density is significantly lower than that of non-BS VIP neurons (Supplementary Figure 2). Roman numerals are layers, dashed lines layer borders.

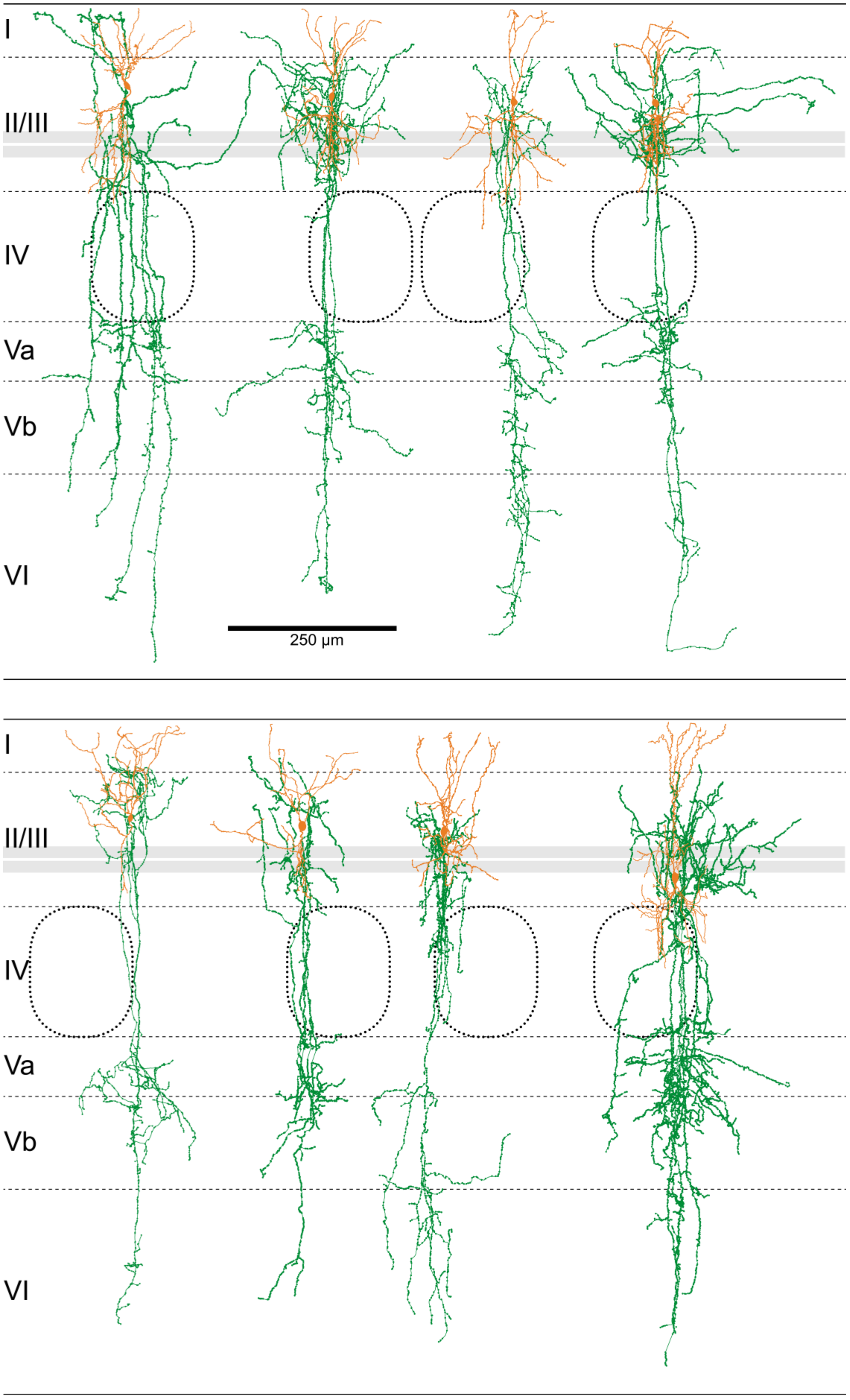

I
II/III
IV
Va
Vb
VI
250 µm
I
II/III
IV
Va
Vb
VI

Supplementary Figure 6: Reconstruction of the morphology of 8 individual non-BS VIP neurons

The 8 individual BS VIP neurons are shown with somata and dendrites in orange and axon in green with their corresponding home barrel in layer IV. Grey rectangles indicate the area in which their dendritic density is significantly lower than that of non-BS VIP neurons (Supplementary Figure 2). Roman numerals are layers, dashed lines layer borders.

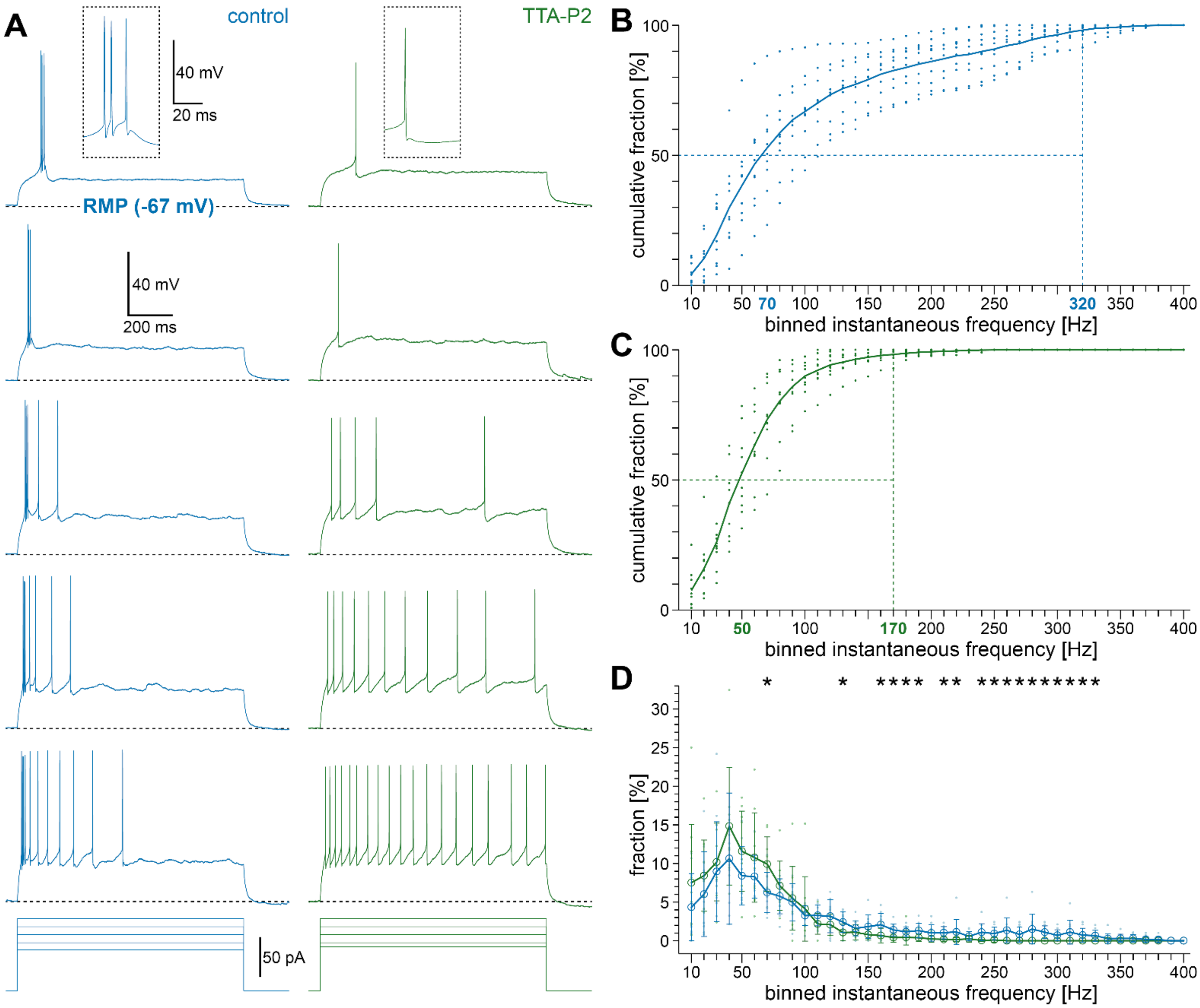

A
control
TTA-P2
40 mV
20 ms
RMP (-67 mV)
40 mV
200 ms
50 pA
B
cumulative fraction [%]
binned instantaneous frequency [Hz]
70
320
C
cumulative fraction [%]
binned instantaneous frequency [Hz]
50
170
D
fraction [%]
binned instantaneous frequency [Hz]

Supplementary Figure 7: T-type calcium channels mediate burst firing in BS VIP neurons

(A) Responses of a single BS VIP neuron at RMP under control conditions (-67 mV; blue) and during bath-application of TTA-P2, a T-type calcium channel antagonist (green). Top traces depict action potentials at rheobase stimulation (insets show APs with a higher temporal resolution), followed by responses to increasing stimulation strength (stimulating rectangular pulses beneath recordings). (B) Frequency spectrum of 10 BS VIP neurons at RMP under control conditions. 98% of all ISIs are found in a frequency range from 10 to 320 Hz (blue stippled vertical line, solid line is mean per bin, dots indicate individual data points) and the half-maximum reached at 70 Hz (stippled horizontal line). (C) Frequency spectrum of BS VIP neurons at a membrane potential of -50 mV (plot as in B). Under TTA-P2, 98% of all ISIs are found in a frequency range from 10 to 170 Hz and half-maximum reached at 50 Hz. (D) Comparison of the frequency spectra in burst and tonic mode. Under TTA-P2, the fraction of IFFs significantly increases in the frequency bin of 70 Hz (green; open circles mark average per bin; error bars SD) in comparison to control conditions (blue). In the high frequency range above 130 Hz, corresponding to IFFs within bursts, the fraction of IFFs significantly decreases in several bins due to the absence of bursts when T-type calcium channels are blocked.

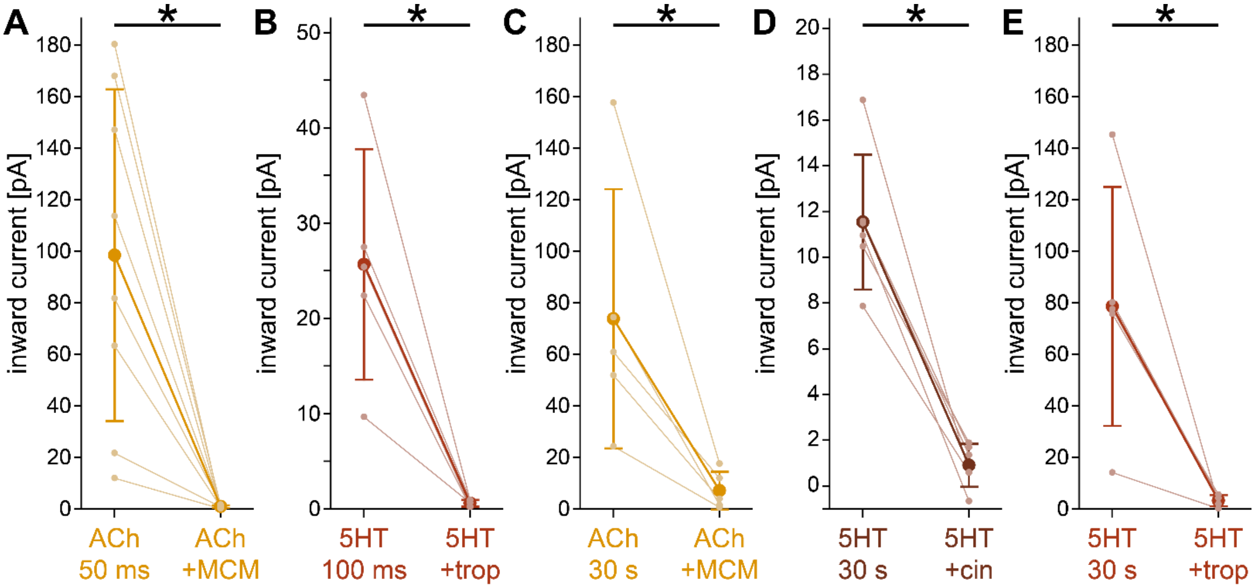


Supplementary Figure 8: Quantification of the effects of antagonists against nicotinic AChR, $5HT_{3a}R$, and $5HT_2R$ in VIP neurons

For single neuron examples see Figure 5. (A) Plot showing that mecamylamine (MCM, 100 µM), a nicotinic AChR antagonist, blocked inward currents evoked by 50 ms of focal pressure application of ACh (100 µM) in VIP neurons (n = 8; large circles and dark line depict average, error bars SD, small circles individual data points connected by lines). (B) Tropisetron (trop, 10 nM), a $5HT_{3a}R$ antagonist, blocked currents evoked by 100 ms of focal pressure applications of 5HT (200 µM) in VIP neurons displaying a response to 5HT under these conditions (n = 5). (C) Mecamylamine (100 µM) also blocked currents evoked by 30 s of focal pressure application of ACh (100 µM, n = 5). (D) Cinanserin (cin, 400 µM), a $5HT_2R$ antagonist, blocked currents evoked following 30 s focal pressure application of 5HT (200 µM, n = 6). (E) Tropisetron blocked currents evoked during 30 s of focal application of 5HT (200 µM, n = 5).

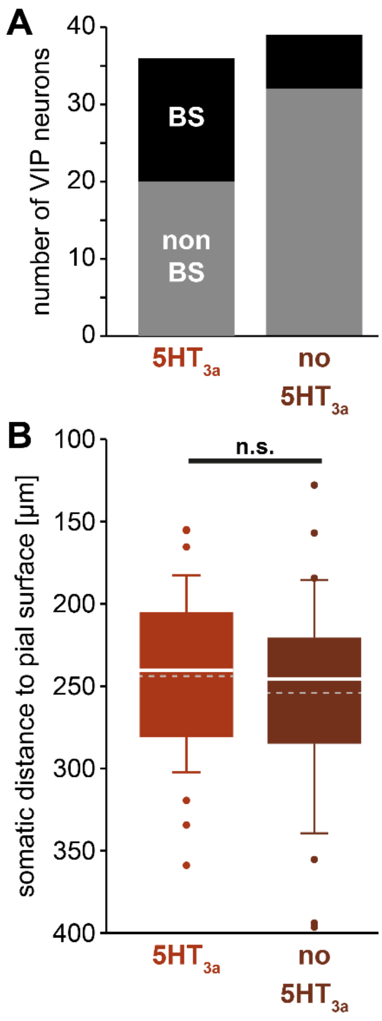


Supplementary Figure 9: Quantification of 5HTR3a activity in VIP neurons

(A) Bar graph depicting the number of VIP neurons displaying 5HT3aR-mediated responses (36 of 75; left bar) and no 5HT3aR-mediated responses (39 of 75, right bar) as well as the number of BS VIP neurons in each group (black). (B) Box plots showing that somata of VIP neurons with 5HTR3a-mediated responses (left; light red) were distributed throughout layer II/III in a similar manner as those without (right; dark red; dashed lines depict average, solid lines mean).

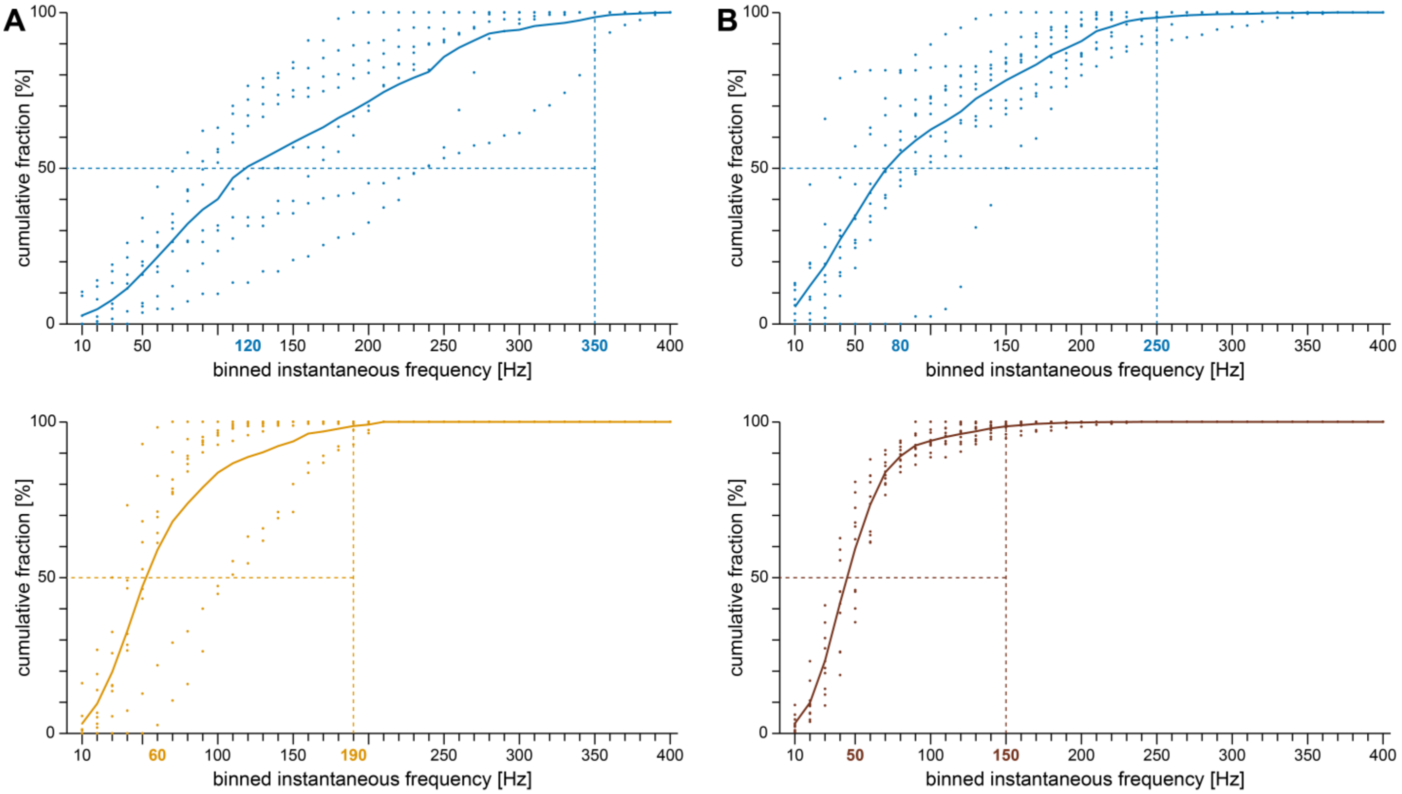


Supplementary Figure 10: Cumulative frequency plots under control conditions and during neuromodulation by ACh or 5HT

Analysis of IFFs of those BS VIP neurons used for bath-application of ACh or 5HT. IFFs of individual BS VIP neurons were pooled, quantified in 10 Hz bins, and relative proportions plotted cumulatively. Stippled vertical lines indicate the frequency bin corresponding to 98% of the maximum, stippled horizontal lines indicate the half-maximal proportion. Solid lines are mean per bin, dots indicate individual data points. The frequency bins at which the half-maximal proportion as well as 98% of the maximum are reached are highlighted by colored numbers. Control recordings were done at RMP, recordings in the presence of neuromodulators were done at depolarized membrane potentials. (A) Frequency spectrum (n = 8) under control condition (upper plot; blue) and during ACh (lower plot; orange). ACh shifted both characteristic frequency features (shown by stippled lines) to lower frequency bins. (B) Frequency spectrum (n = 8) under control condition (upper plot; blue) and during 5HT (lower plot; brown). 5HT also shifted both characteristic frequency features to lower frequency bins.

| | RMP [mV] | | input resistance [MΩ] | | time constant [ms]* | | capacitance RMP [pF] | |
|---|---|---|---|---|---|---|---|---|
| | BS | non-BS | BS | non-BS | BS | non-BS | BS | non-BS |
| n | 55 | 214 | 55 | 214 | 55 | 214 | 55 | 214 |
| Mean | -65.1 | -66.3 | 337.2 | 294.2 | 18.5 | 15.5 | 51.2 | 48.4 |
| SD | 3.5 | 3.6 | 112.9 | 110.9 | 5.2 | 5.4 | 10.3 | 12.5 |
| Median | -65.2 | -66.6 | 315.8 | 271.0 | 17.5 | 14.4 | 50.0 | 46.0 |
| 25% | -67.8 | -68.6 | 241.4 | 211.8 | 15.3 | 11.9 | 45.1 | 40.0 |
| 75% | -62.4 | -63.6 | 412.4 | 348.7 | 21.7 | 18.3 | 57.7 | 53.6 |
| P | 0.735 | | 0.09 | | <0.001 | | 0.315 | |

| | sag [%]* | | rectification index [%] | | rheobase [pA]* | | AP latency [ms]* | |
|---|---|---|---|---|---|---|---|---|
| | BS | non-BS | BS | non-BS | BS | non-BS | BS | non-BS |
| n | 55 | 214 | 55 | 214 | 55 | 214 | 55 | 214 |
| Mean | 12.4 | 9.5 | 12.8 | 13.6 | 51.6 | 73.2 | 113.0 | 92.8 |
| SD | 5.5 | 5.3 | 1.9 | 2.7 | 22.0 | 36.6 | 41.5 | 56.5 |
| Median | 12.0 | 8.8 | 12.5 | 13.1 | 49.1 | 67.9 | 111.3 | 78.8 |
| 25% | 7.7 | 5.5 | 11.4 | 11.8 | 36.1 | 46.9 | 85.3 | 49.9 |
| 75% | 16.1 | 12.3 | 14.3 | 14.7 | 60.2 | 94.4 | 135.9 | 117.3 |
| P | <0.001 | | 1.755 | | <0.001 | | <0.001 | |

| | AP FT [mV] | | AP amplitude [mV] | | AP time to peak [ms]* | | AP width [ms]* | |
|---|---|---|---|---|---|---|---|---|
| | BS | non-BS | BS | non-BS | BS | non-BS | BS | non-BS |
| n | 55 | 214 | 55 | 214 | 55 | 214 | 55 | 214 |
| Mean | -35.4 | -35.0 | 66.9 | 68.6 | 0.367 | 0.402 | 0.448 | 0.524 |
| SD | 2.7 | 2.9 | 8.2 | 7.9 | 0.054 | 0.060 | 0.097 | 0.106 |
| Median | -35.4 | -35.1 | 66.2 | 69.5 | 0.351 | 0.395 | 0.416 | 0.525 |
| 25% | -37.6 | -37.2 | 62.3 | 63.5 | 0.326 | 0.357 | 0.370 | 0.445 |
| 75% | -33.8 | -32.9 | 73.4 | 74.2 | 0.393 | 0.440 | 0.495 | 0.602 |
| P | 3.375 | | 2.115 | | <0.001 | | <0.001 | |

| | AP slope [V/s] | | AHP peak [mV] | | AHP amplitude [mV] | | AHP time to peak [ms]* | |
|---|---|---|---|---|---|---|---|---|
| | BS | non-BS | BS | non-BS | BS | non-BS | BS | non-BS |
| n | 55 | 214 | 55 | 190 | 55 | 190 | 55 | 190 |
| Mean | 240.9 | 227.2 | -45.5 | -46.2 | 10.1 | 11.1 | 0.497 | 0.718 |
| SD | 45.2 | 44.4 | 3.5 | 3.5 | 2.8 | 3.1 | 0.127 | 0.246 |
| Median | 242.6 | 228.9 | -45.4 | -46.0 | 10.0 | 11.0 | 0.493 | 0.680 |
| 25% | 211.0 | 198.1 | -47.9 | -48.7 | 12.3 | 12.8 | 0.401 | 0.535 |
| 75% | 271.5 | 259.0 | -43.4 | -43.8 | 7.8 | 9.2 | 0.573 | 0.882 |
| P | 0.633 | | 2.91 | | 1.17 | | <0.001 | |

Supplementary Table 1: Comparison of 16 electrophysiological properties between BS and non-BS VIP neurons

Statistically significant differences are marked in grey. RMP = resting membrane potential; R in = input resistance; FT = firing threshold; SD = standard deviation. T-tests were performed if data passed a normality test (Shapiro-Wilk), Mann-Whitney rank sum tests if data failed a normality test, P-Values were corrected post hoc to preclude alpha inflation (Bonferroni correction).

| | RMP [mV]* | | rheobase [pA]* | | AP latency [ms] | | AP FT [mV] | |
|---|---|---|---|---|---|---|---|---|
| | burst | tonic | burst | tonic | burst | tonic | burst | tonic |
| n | 50 | 49 | 50 | 49 | 50 | 49 | 50 | 49 |
| mean | -65.2 | -50.8 | 49.6 | 19.1 | 114.1 | 169.0 | -35.5 | -34.4 |
| SD | 3.5 | 1.5 | 19.5 | 21.1 | 42.6 | 128.7 | 2.6 | 3.2 |
| median | -65.3 | -50.8 | 49.0 | 14.1 | 111.3 | 137.4 | -35.5 | -34.5 |
| 25% | -67.9 | -51.4 | 36.2 | 6.8 | 85.5 | 80.6 | -37.6 | -36.7 |
| 75% | -62.7 | -49.6 | 60.0 | 25.6 | 137.3 | 221.0 | -33.8 | -32.0 |
| P | <0.001 | | <0.001 | | 0.67 | | 0.52 | |

| | AP amplitude [mV] | | AP time to peak [ms]* | | AP width [ms] | | AP slope [V/s]* | |
|---|---|---|---|---|---|---|---|---|
| | burst | tonic | burst | tonic | burst | tonic | burst | tonic |
| n | 50 | 49 | 50 | 49 | 50 | 49 | 50 | 49 |
| mean | 67.9 | 64.8 | 0.364 | 0.383 | 0.446 | 0.466 | 245.7 | 221.1 |
| SD | 7.4 | 8.0 | 0.051 | 0.465 | 0.095 | 0.089 | 41.2 | 34.7 |
| median | 67.4 | 65.4 | 0.350 | 0.378 | 0.414 | 0.447 | 245.8 | 218.4 |
| 25% | 63.0 | 58.8 | 0.325 | 0.343 | 0.369 | 0.401 | 211.9 | 196.7 |
| 75% | 73.7 | 71.3 | 0.393 | 0.409 | 0.497 | 0.516 | 277.4 | 238.7 |
| P | 0.47 | | 0.013 | | 1.16 | | 0.02 | |

| | AHP peak [mV] | | AHP amplitude [mV]* | | AHP time to peak [ms]* | |
|---|---|---|---|---|---|---|
| | burst | tonic | burst | tonic | burst | tonic |
| n | 50 | 47 | 50 | 47 | 48 | 42 |
| mean | -45.6 | -46.4 | 10.1 | 12.2 | 0.478 | 0.644 |
| SD | 3.4 | 3.7 | 2.8 | 3.1 | 0.098 | 0.130 |
| median | -45.4 | -46.2 | 10.3 | 12.3 | 0.477 | 0.665 |
| 25% | -47.9 | -49.4 | 12.3 | 14.8 | 0.399 | 0.532 |
| 75% | -43.8 | -43.9 | 7.8 | 9.7 | 0.550 | 0.743 |
| P | 1.53 | | <0.001 | | <0.001 | |

Supplementary Table 2: Comparison of 11 BS VIP neuron electrophysiological properties between burst and tonic mode

Statistically significant differences are marked in grey. RMP = resting membrane potential; FT = firing threshold; SD = standard deviation. T-tests were performed if data passed a normality test (Shapiro-Wilk), Mann-Whitney rank sum tests if data failed a normality test.
